# A novel bioinformatics approach to reveal the role of circadian oscillations in AD development

**DOI:** 10.1101/2020.07.16.207357

**Authors:** Zhiwei Ji, Dazhi Shang, Pora Kim, Mengyuan Yang, Sijia Wu, Weiling Zhao, Eunju Kim, Marvin Wirianto, Zheng Chen, Seung-Hee Yoo, Xiaobo Zhou

## Abstract

Altered circadian gene expression may contribute to Alzheimer’s disease (AD) progression. Unfortunately, sampling the central nervous system (CNS) at multiple time points is not feasible. Moreover, there are no AD-related time-series transcriptome datasets available for studying these circadian patterns and their impacts on AD development. In this study, we introduce a novel computational platform, Event-driven Sample Ordering for Circadian Variation Detection (ESOCVD), to reveal rhythmic patterns of gene expression of AD using untimed transcriptome datasets. ESOCVD was applied to 20 untimed gene expression profiles of 16 brain regions from approximately 3000 AD patients in public transcriptome databases. Our analysis revealed five types of circadian alteration patterns in ~2,000 circadian genes in different brain regions of AD patients. Further analyses of additional databases confirmed that our analytical platform can be applied to identify the evolutionary dynamics of circadian variation during the process of AD development. Through the gene expression correlation analysis for our 8 circadian genes identified from AMP-AD MSBB cohorts, we identified stage-specifically enriched biological processes with anticipated context. Gene expression analysis of AD mouse brain tissues further substantiated the predictions of the ESOCVD model. In summary, ESOCVD is highly versatile in bridging circadian research and precision medicine.

## Introduction

The circadian clock is a complex biological machine that allows organisms, from cyanobacteria, fruit flies to humans, to predict and prepare for the challenges of daily life^1^. The body’s clock system maintains 24-h rhythms in physiological functions, including the sleep-wake cycle, and synchronizes them to the light-dark cycle^2^. The clock system is composed of cell-autonomous molecular oscillators^3,4^, and the molecular oscillator, in turn, consists of conserved transcriptional and translational feedback loops of ‘core clock genes’ ^5,6^. These clock genes drive circadian oscillations of so-called clock-controlled genes (CCGs) at the cellular level to regulate various cellular and systemic functions^7,8^. Disturbances of the circadian rhythms have long been associated with many neurological and psychiatric diseases^9,10^, including Alzheimer’s disease (AD)^1,11^, Parkinson’s disease (PD)^12,13^, and Huntington’s disease (HD)^14,15^. Therefore, mechanistic understanding of the interactions between circadian biology and neurodegenerative pathophysiology may provide new therapeutic opportunities against neurodegenerative diseases.

Alzheimer’s disease affects 20-40 million people worldwide and is the most common cause of progressive cognitive dysfunction in adults. Circadian dysfunction phenotypes are observed at the early stage of AD^1^. Drosophila with overexpressed amyloid-β (Aβ) peptides also show a severely disturbed circadian function^16^. AD mouse models show accelerated disintegration of circadian rhythms with age as compared to normal counterparts^17–21^. Several studies in transgenic mice demonstrate that amyloid deposition in the brain leads to the disruption of normal sleep architecture. Abnormalities in mice include a loss of robust day-night oscillation in brain lactate and altered nocturnal activity level^17,20,21^. Furthermore, ~80% of AD patients over 65 years old suffer from circadian rhythm disorders, such as disturbances in thermoregulation and sleep-wake cycles^22,23^. AD patients exhibit disturbances in the timing and duration of the sleep cycle, primarily manifested as increased wakefulness at night and increased sleep during the day, which can progress to a loss of day-night variation^11^.

Although circadian dysfunction has long been considered as a prevailing symptom of AD, it has more recently been hypothesized as a contributor to the pathogenesis of AD^24–26^. New findings in the fruit fly and rodent models indicate that the deletion of the clock gene *Bmal1, Clock, Per1,* and *Cry1* may cause an accelerated aging phenotype characterized by an earlier decline in cognitive functions^5,25,27^. Recent experiments found that sleep deprivation caused a striking increase in the Aβ plaque burden in mice that express AD-associated mutant forms of human amyloid precursor protein *(APP)* and develop *Aβ* plaques with age^28,29^. A subsequent study using mice that express human *APP*, *PS1,* and human *Tau* transgenes also demonstrated an increase in cortical *Aβ* plaque burden following chronic sleep deprivation^30^. Moreover, human data also support a role for circadian disruption in AD pathogenesis. An epidemiologic study of daily activity from over 1,200 initially cognitively-normal older women demonstrated that diminished circadian rhythm amplitude, robustness, or phase delay were associated with increased risk of developing dementia during the 5-year follow-up period^31^. However, it remains unclear whether or how the circadian system influences AD pathogenesis. Therefore, a better understanding of how circadian dysfunction contributes to neurodegenerative disease mechanisms will facilitate the development of novel therapies for AD.

The abnormalities in circadian-ordered gene expression are linked to an array of downstream pathobiology of AD, including oxidative stress, misfolded proteins, and epigenetic burden^32,33^. Identifying the dysregulation in the oscillatory gene expression patterns from the existing transcriptome datasets will provide novel mechanistic insights into the complex etiology of AD. Unfortunately, there are few public transcriptome datasets with multiple “timed” control human samples to study such patterns, and none available for AD. However, it has been shown that compiling a collection of single “untimed” gene expression samples can generate a snapshot of circadian gene expression. This can be achieved by analyzing the existing large transcriptome datasets with computational approaches/models.

In recent years, several computational algorithms have been developed to detect oscillatory patterns of gene expression in health and disease, e.g., supervised learning (ZeitZeiger^34^, JTK_CYCLE^35^, TimeSignature ^36^, *etc*.) and unsupervised learning (Oscope^37^, and CYCLOPS^38^, *etc*.). The limitation of the first three approaches is that they only work on timed gene expression data. Oscope is designed to extract the oscillatory dynamics of gene groups from single-cell RNA-seq data. However, it requires much more computation and is sensitive to the ordering of cells. CYCLOPS can estimate the circadian phase of each sample with untimed gene expression profiles. However, the requirement for a large sample population to fill the entire periodic cycle is the key limitation. Therefore, developing efficient, robust, and data-insensitive algorithms for circadian variation detection on untimed transcriptomic datasets remains a big challenge.

In this study, we proposed a computational approach, Event-driven Sample Ordering for Circadian Variation Detection (ESOCVD), to decipher the oscillatory patterns of gene expression in AD using “untimed” transcriptome datasets. ESOCVD extracts the global pacemakers of the clock system to predict the circadian phase of each sample by modeling time-dependent events with the Bayesian approach. Based on the inferred circadian phases, we tracked five types of circadian variation in gene expression during AD progression. To verify the effectiveness, we applied the ESOCVD model to 20 gene expression profiles collected from public databases AMP-AD^39^, and GEO (Gene Expression Omnibus), which cover 16 brain regions of ~3000 AD patients. We mainly focused on 1691 seed genes and 857 potential brain-related circadian genes (PBCG) (Supplementary Fig. 1). The outcomes of ESOCVD revealed the heterogeneity of the occurrence of circadian variations in different brain regions of AD patients. Functional analysis indicates that there are five regions associated with circadian rhythm regulation, including superior frontal gyrus (SFG), superior temporal gyrus (STG), *etc*. The estimated circadian rhythms of well-known clock genes and circadian genes under normal conditions are consistent with these reported in Circadian Gene Database (CGDB)^40^. Moreover, we checked 21 AD-related clock-controlled genes^25,41^ and found that loss or gain of rhythmicity in multiple-regions. In addition, we developed a systems biology approach to reconstruct gene regulatory network (GRN) connecting AD development and circadian rhythm processes. We successfully identified three signaling pathways for brain-region SFG, Visual cortex (VC), and STG, respectively. Finally, we tracked the evolution of the cycling rhythm of circadian genes during the process of AD development. Through studying 8 circadian genes in AMP-AD MSBB cohorts^42^, we identified 16 AD marker SNPs and 85 correlated genes that were significantly associated with these genes in disease stages. These genes showed stage-specifically enriched biological processes with the anticipated context of disease stages. More data sources provide evidence that 8 circadian genes act as intermediators in AD pathogenesis.

## Results

### The datasets for model validation

In this study, we collected untimed transcriptomic data from existing datasets to validate our ESOCVD approach. These gene expression profiles are related to 16 regions of the brain from 2930 subjects with or without dementia (Supplementary Table 1). The samples from GSE125583, GSE5281, GSE15222, GSE44772, and GSE29378 were labeled as control and AD. Both MSBB and GSE131617 include patients from early to late stage of AD. All the subjects involved in GSE131617 were scored using Braak staging^43^.

We defined the samples with stage “0” as control, “1-2” as early, “3-4” as mid, and “5-6” as late. The samples in MSBB are grouped based on cognitive status (Clinical Dementia Rating scores, CDR^44^). Thus, we assigned the samples with CDR score “0” as control, “0.5” and “1” as early, “2” as mid, and “3-5” as late. Here, ESOCVD was used to detect the circadian alteration in any stage of AD relative to control.

### Eatimatation of circadian phases using ESOCVD

For the time series transcriptome, circadian patterns can be easily detected according to the series of sampling time^35,45–47^. However, the order of samples in an untimed dataset is unknown. ESOCVD optimally combines eigengene patterns to reveal an underlying circadian phase of each sample in a single cycle, and then uses the inferred phases to order the data (Fig. 1). Each eigengene is defined as an oscillating biomarker and evolves following a circadian pattern (see the details in “**Methods**”). Furthermore, the expected expression distribution of an eigengene can be determined based on the relative positions of ‘crest’ and ‘tough’ in a single cycle. Therefore, we defined an event sequence **S**, denoting the occurrence of the ‘crest’ and ‘tough’ (*z*_*i*1_ or *z*_*i*2_) for all the eigengenes. ESOCVD quickly searches the optimal sequence **S** by maximizing the likelihood of the expression data. In this study, we chose the number of eigengenes (indexed as *i* = 1, …,*I*) is 12 and the distance between any pair of *z*_*i*1_ and *z*_*i*2_ is half cycle (12 hours). In other word, each event in **S** denotes 1 hour. The 24 events separate the time axis to 25 progression stages. After maximizing the likelihood in **Eq. (1)**, we obtained the optimal sequence **S**. In the meantime, we got a *N* *25 matrix *pk*, which represents the probability of each subject *j* at each stage (1 ≤ *j ≤ N*). The subject *j* is assigned to stage *k* (0 ≤ *k* ≤ 24) with maximal probability (**Eq. (2)**). Once all the samples were assigned to corresponding phases in a single cycle, the circadian rhythmicity can be easily detected by using Cosinor model^48^.

**Fig. 1.**
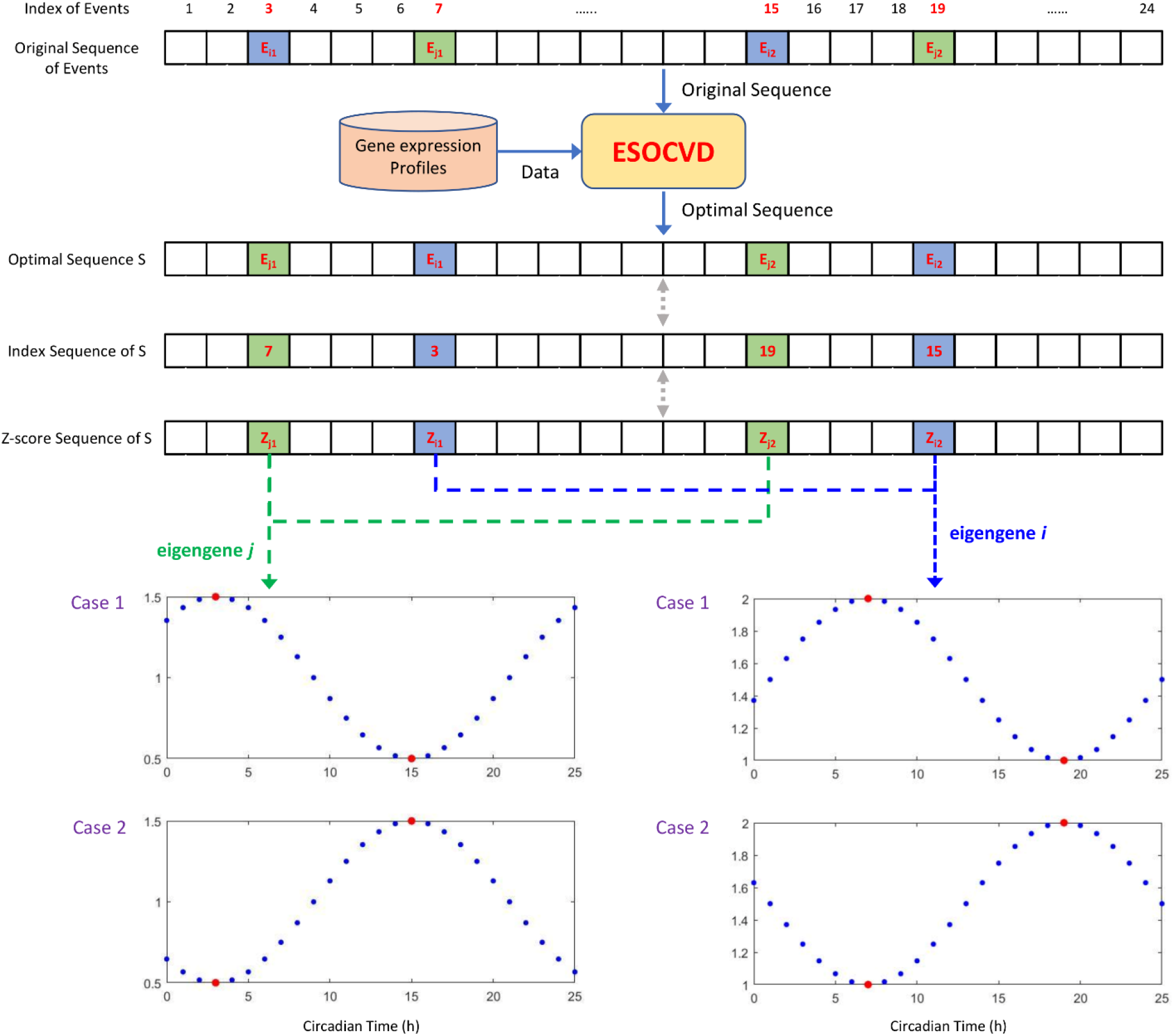
The flowchart of ESOCVD model.

We examined GSE125583 as an example. Supplementary Fig. 2 presents the inferred distribution of all the samples across the stages. We found that control samples distributed evenly in a single cycle. However, AD samples mainly distributed in the 1^st^ and 4^th^ quarters. Similarly, the circadian phases of all the samples associated with each data profile were estimated (Supplementary File1).

### Detection of the circadian variations of gene expression

Based on the inferred order of samples, we detected the circadian variations of gene expression between control and AD groups with cosinor regression^48^. ESOCVD assigns each sample a phase (stage) along a single cycle (0 ≤ 0 ≤ 24Λ). Given the ESOCVD-ordered expression data of control or AD samples, we used cosinor regression to fit the expressions of each gene in each group and identified the genes that have significantly different rhythmic patterns between AD and control samples (P-value<0.05).

Then, we examined five types of circadian variations across the genome: 1) amplitude change; 2) loss of rhythmicity (LR); 3) gain of rhythmicity (GR); 4) Phase shift; 5) Base shift (see the details in “**Methods**”). For each data profile, we mainly focused on two groups of genes (Supplementary Fig. 1) and then examined the gene expression with circadian variations. The first group has 1691 seed genes, including 1670 circadian genes collected from mouse, and 21 AD-related clock-controlled genes. In addition, we searched GO database^49^ and selected 15 core clock genes, which are also included in the seed gene list. The second group includes 631 brain-related circadian genes reported in CGDB^40^, 288 aging-related circadian genes^45–47,50,51^, and 34 AD risk factors. Supplementary Fig. 1 shows that 700 PBCG genes do not belong to the seed gene list. The details of seed genes and PBCG genes were described in Supplementary File2.

Fig. 2 shows the numbers of seed genes with circadian variation predicted by ESOCVD in 16 brain regions. We found that loss or gain of rhythmicity occurred in multiple regions, while amplitude change and phase shift only appear in the 1 or 2 areas. There are more genes lose the circadian rhythm in SFG, STG, and DPC (dorsolateral prefrontal cortex), *etc*. In Fig. 2a, the first 16 columns are associated with the AD patients from United States, while the last 3 columns (marked with “*”) are relevant to Japanese patients. It seems that there are more LR genes and fewer GR genes in FC (Frontal cortex) and EC (Entorhinal cortex) regions of Japanese patients, which is opposite to the situations on American patients. Moreover, Fig. 2b shows the periodic rhythms of 10 core clock genes in multiple brain regions of control samples. We found that the circadian phases of “peak” and “trough” for all the clock genes are very close to the evidence reported in the database CGDB, which indicates that our approach is reliable (Supplementary Table 2).

**Fig. 2.**
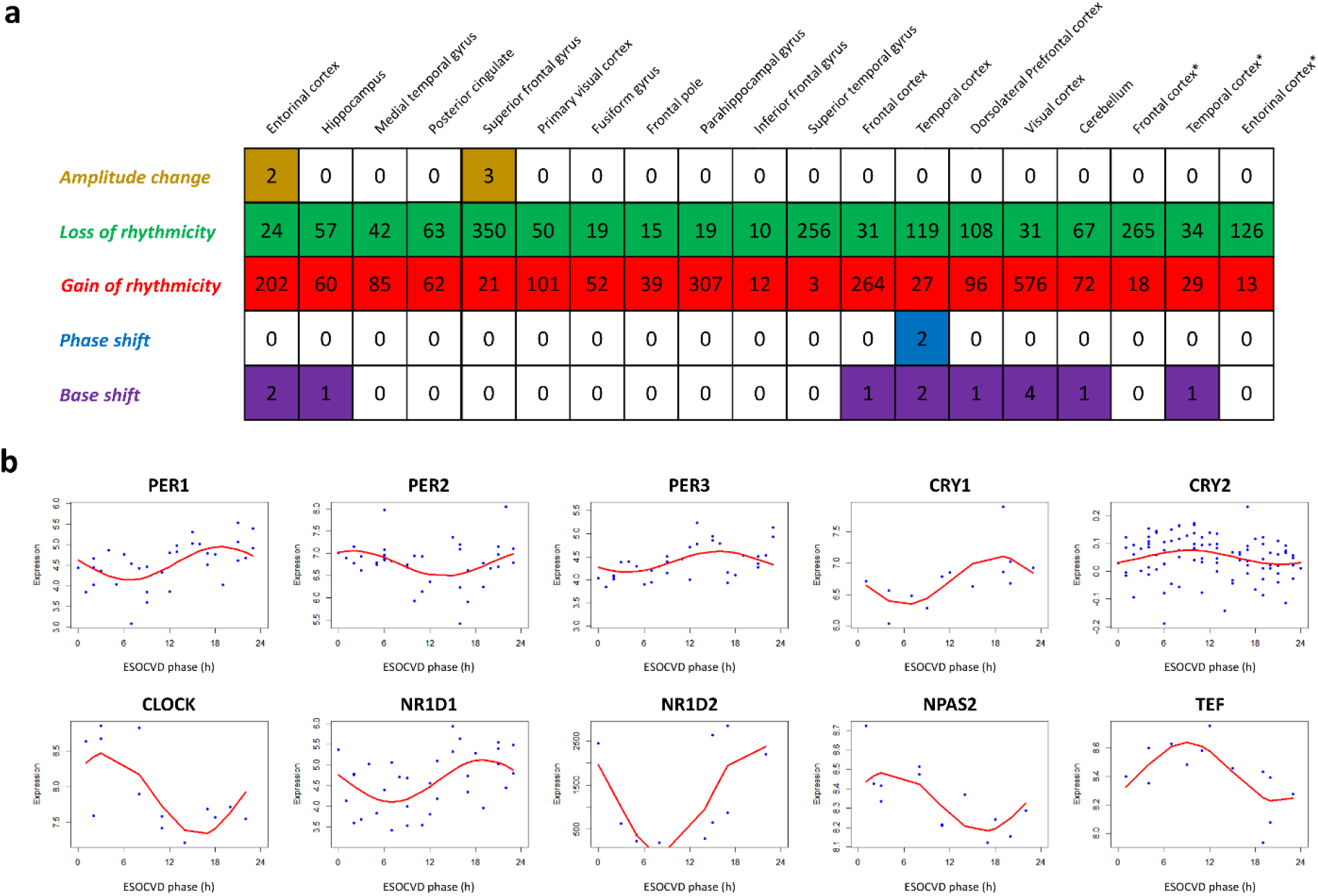
ESOCVD analysis of circadian transcriptome in late on-set of AD. **a,** the number of seed genes with circadian changes in multiple brain regions detected by ESOCVD. **b,** Reconstructed expression profiles of selected clock genes are plotted as a function of ESOCVD phase. Each dot represents one sample in control group.

We further explored the circadian patterns of 21 AD-related clock-controlled genes across all the brain regions (Fig. 3a). Eighteen genes show circadian alterations in some brain regions, while 3 genes displayed no changes. Most of them occurred LR or GR in different regions (Supplementary Table 3). Fig. 3b presents 6 selected AD-related clock-controlled genes, which are lost the rhythmicity in a particular region. Moreover, we performed qPCR experiments using cortex from APP/PS1 model mice to validate the cycling patterns of key circadian genes identified by ESOCVD. Fig. 3c indicates that the observed trends of diurnal expression of these genes in wide-type (WT) mice are highly consistent with our prediction of the control group, and the circadian rhythm detected in WT is markedly diminished in the AD group.

**Fig. 3.**
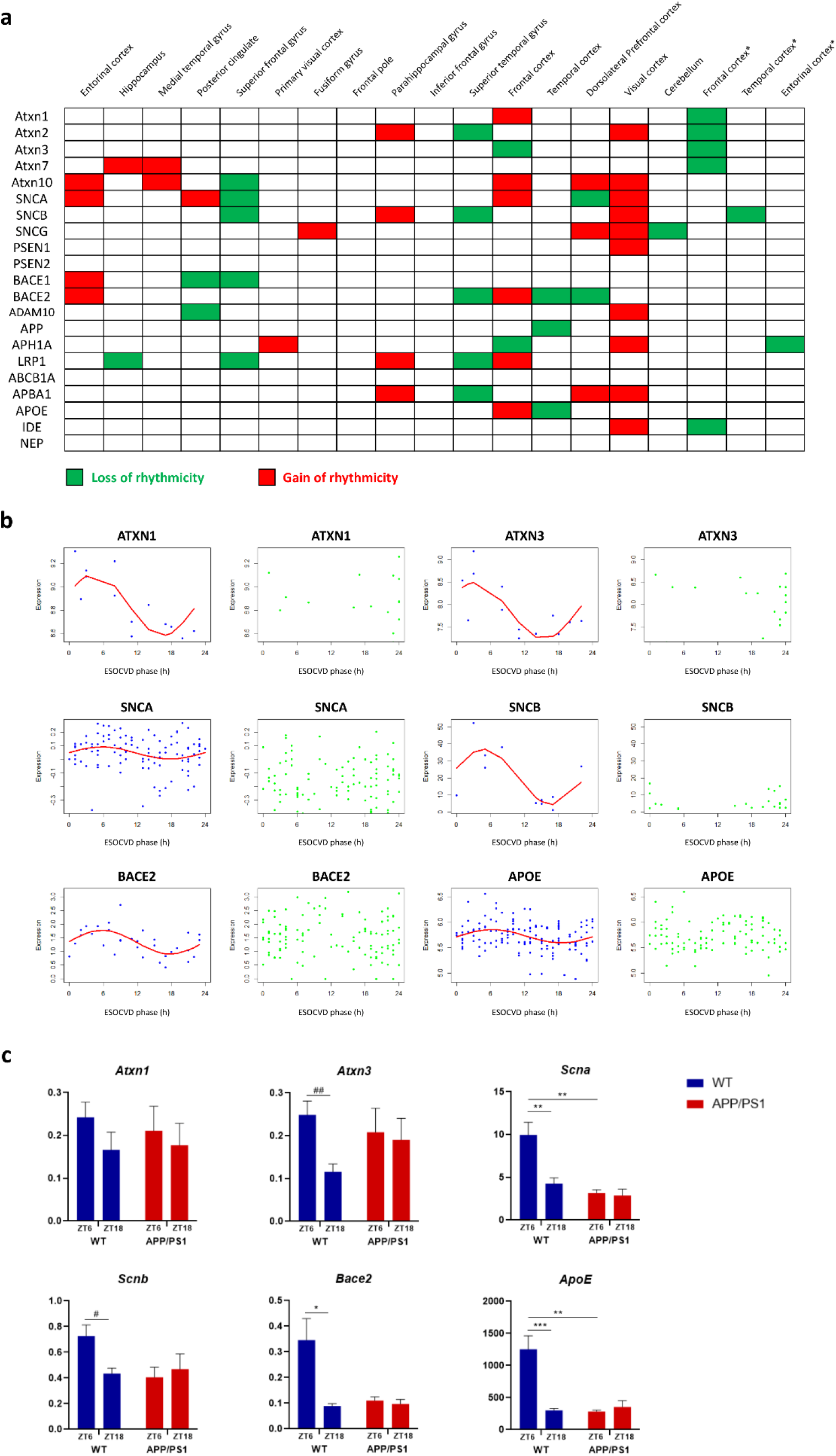
ESOCVD analysis of AD-related circadian changes in late on set of AD. **a,** the distribution of circadian changes of AD-related clock-controlled genes in multiple brain regions detected by ESOCVD. **b,** Reconstructed expression profiles of six AD-related circadian genes are plotted as a function of ESOCVD phase. ATXN1 and ATX3 are plotted with GSE131617_FC. SNCA and SNCB are related with DPC (GSE44772) and SFG (GSE5281). BACE2 and APOE are plotted with MSBB_STG and GSE15222_TC, respectively. The dots with blue and green color represent control and AD samples, respectively. **c,** Real-time qPCR analysis of six genes presented in **(b)** in the cortex. Error bars represent SEM (n=4-6) # p<0.05, ## p<0.01, unpaired t-test; *p<0.05, **p<0.01, ***p<0.001,two-way ANOVA with Sidak multiple comparisons test.

Similarly, we summarized the circadian changes of 700 PBCG genes, as shown in Fig. 4a. The distributions of LR and GR are very similar to that in seed genes (Fig. 4a, Fig. 2a). The other three types of patterns appear in only 2-3 regions. Fig. 4b shows 15 selected circadian genes detected by ESOCVD, which are representative samples to elucidate five types of patterns. The genes *VEGFD, ALKBH3,* and *MRPL41* in the first row showed amplitude change in the AD group. Particularly, the angiogenesis mediator *VEGFD* has a twofold reduction in amplitude in AD samples, which is consistent with Bridel’s findings that *VEGFD* reveals a tendency towards a decrease with AD severity^52^. *PSMD5, KBTBD11,* and *SDR39U1* lost rhythmicity in AD, hinting at previous findings that deletion/disruption of these genes may cause behavioral abnormalities in human^53,54^. Moreover, *ENO3*, *SMAD5*, and *SUGT1* gained rhythmicity in AD samples relative to controls, demonstrating the ability of *ENO3* to rescue neuronal loss^55^. In addition, *DNAJB5, SMPD4,* and *ATP13A5* exhibited a clear phase shift in the AD group compared with the control group. Finally, *OMG, FRMD4A,* and *CYTB* displayed baseline shift, indicating that the mean values of gene expression are significantly reduced in AD samples. The details of all the genes with circadian variation are described in Supplementary File 3.

**Fig. 4.**
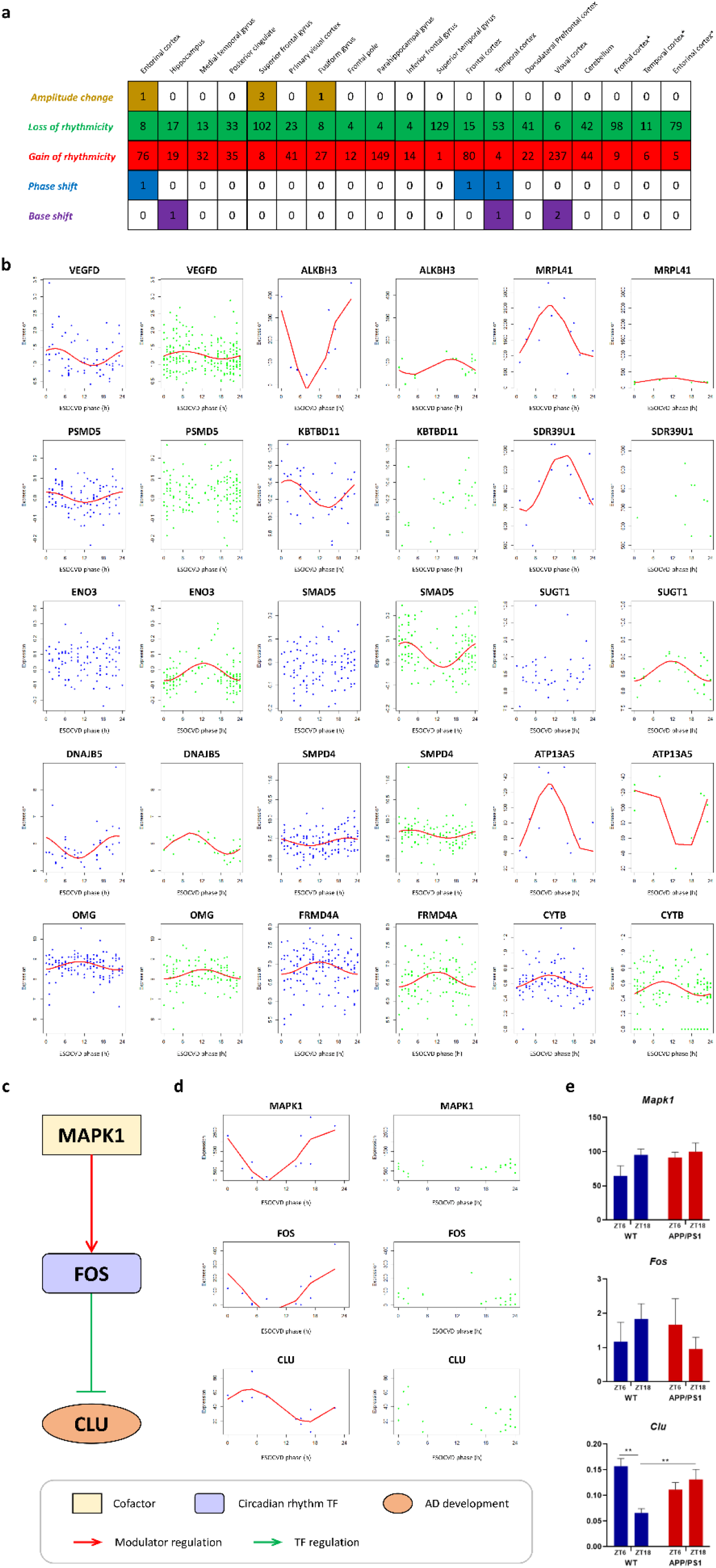
ESOCVD analysis of PBCG-related circadian changes in late on set of AD. **a,** the number of PBCG genes with circadian changes in multiple brain regions detected by ESOCVD. **b,** Reconstructed expression profiles of selected PBCG genes are plotted as a function of ESOCVD phase. **c,** Inferred CLU-related signaling pathways in SFG. **d,** Reconstructed expression profiles of genes on CLU pathways are plotted as a function of ESOCVD phase. **e**, qPCR experiments for CLU pathway in the cortex. Error bars represent SEM (n=4-6). *p<0.05, **p<0.01, ***p<0.001,two-way ANOVA with Sidak multiple comparisons test. The dots in (b) and (d) with blue and green color represent control and AD samples, respectively.

### Pathway analysis of circadian changes in AD samples

After obtaining the signatures with the circadian changes, we conducted pathway-level analysis using bioinformatics platform DAVID^56^ to assess whether these inferred genes are associated with important pathways or biological processes. The top 5 significant brain regions include superior frontal gyrus (SFG), superior temporal gyrus (STG), frontal cortex (FC), temporal cortex (TC), and posterior cingulate (PC). KEGG and GO biological process sets that were overrepresented among those genes that change rhythmicity in AD patients are collected, e.g. microglial cell activation^57,58^, circadian rhythm, circadian regulation of gene expression, neuron apoptotic process, and insulin signaling pathways^59,60^, *etc.* The details of DAVID analysis are shown in Supplementary Tables 4-8.

### Exploration of the associations between AD development and circadian rhythm at the network level

Since the molecular mechanism of how circadian dysfunction links AD development remains poorly defined, we developed a systems biology approach to reveal the association between AD development and circadian variation at the network level (Supplementary Fig. 3). Our approach fully utilizes the upstream (cofactor) and downstream (target gene, TG) regulation information around transcription factor (TF) and integrates the functional linkage data and gene expression. By systematically mining the consistency of regulatory relationships, our model infers all the modulatory triplets (cofactor-TF-TG) and constructs gene regulatory network (see the details in “**Methods**”).

Our findings show that 50 AD-risk factors with circadian changes were detected by ESOCVD, e.g., *ABCB1, CLU, BIN1,* and *TREM2, etc.* (Supplementary Table 9). Based on the information for all possible TF-TG pairs in TRRUST^61^, we firstly screened out 2 pairwise transcriptional regulations: 1) *FOS* — | *CLU*^62,63^; 2) *BIN1→MYC^64^*. The analysis on GSE5281 indicates that both *CLU* and its transcription factor *FOS* occur loss of rhythm in SFG (Fig. 4c). *MYC* is the unique target gene of *BIN1^65^.* The transcription factor *BIN1* appears heterogeneity of circadian patterns in VC and STG. Secondly, we identified cofactor-TF regulatory relationships based on the PPI information in STRING^66^. Thirdly, we integrated upstream and downstream regulations of TF in a GRN network. Supplementary Fig. 4 presents the generic GRN pathway maps for *CLU* and *BIN1.* Finally, we inferred the region-specific GRN network by integrating our TILP^67^ approach and gene expression data. Fig. 4c presents the inferred *FOS* pathways connecting circadian rhythm to AD development. The *BIN1*-related pathways are presented in Supplementary Fig. 5. Our analysis shows that the average expressions of *MAPK1* and *FOS* are down-regulated and *CLU* is up-regulated in AD (Supplementary Table 10). Loss of rhythmicity occurred on these 3 genes (Fig. 4d). Our experiment revealed that the observed circadian rhythms on *MAPK1*, *FOS*, and *CLU* are consistent with ESOCVD prediction (Fig. 4e).

Supplementary Fig. 5a presents the inferred *BIN1* pathways associated with VC and STG. *DNM2* expression level was elevated in the AD group and thus activated the downstream regulation through *BIN1* (Supplementary Table 11). The elevated expression of *DNM2* seems to induce the gain of rhythmicity and cause a base increase in downstream. In addition, Supplementary Fig. 5b shows the *BIN1* pathways associated with STG. *BIN2* and *BIN1* were upregulated in AD patients (Supplementary Table 12) and had lose of rhythmicity.

### The evolution of circadian variation during the process of AD development

Since MSBB and GSE131617 include the clinical stage information for each patient, therefore, we explored the dynamic changes in cycling rhythms of seed genes and PBCG genes during the process of AD development. By combining the ESOCVD-inferred cycling rhythm on each condition, we can track the evolution of circadian patterns from early to late-on-set of AD (Fig. 5, Supplementary Fig. 6). Fig. 5a-b show the stage-associated distribution of genes with circadian changes in four brain regions (FP, PG, IFG, and STG) of AD patients in MSBB. Except the superior temporal gyus (STG) has four types of circadian patterns, other three regions only appear LR and GR. We found that the genes with loss of rhythmicity have a higher overlapping rate in three stages, while the genes with gain of rhythmicity have a lower overlapping rate (Supplementary Fig. 7-8). Our data suggests that LR often occurs at an early stage and continues to the late stage, and GR may happen randomly. Similarly, ESOCVD was also applied to the GSE131617 dataset for detecting the evolution of circadian patterns in other three brain regions (FC, TC, and EC). The results are summarized in Supplementary Fig. 6.

**Fig. 5.**
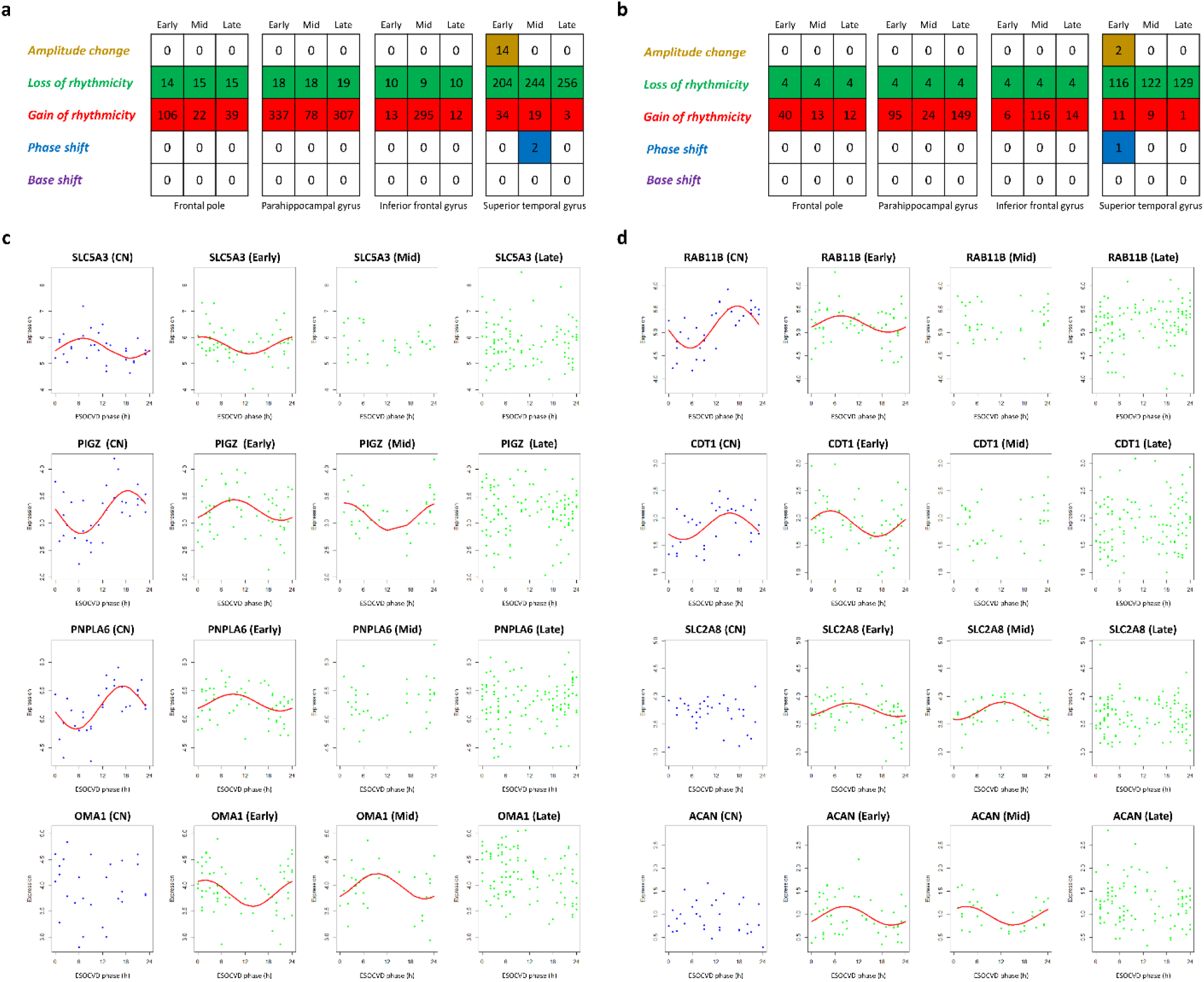
ESOCVD analysis of circadian dynamics in four regions during AD progression. Expression data from MSBB was used. **a,** the number of seed genes with circadian changes in four brain regions detected by ESOCVD. **b,** the number of PBCG genes with circadian changes in four brain regions detected by ESOCVD. **c,** expression of four selected seed genes as a function of ESOCVD phase. PIGZ and PNPLA6 are detected from STG. SLC5A3 and OMA1 are associated with PG and IFG, respectively. **d,** expression of four selected PBCG genes as a function of ESOCVD phase. RAB11B and CDT1 are detected from STG. SLC2A8 and ACAN are associated with FP and PG, respectively. The dots with blue and green color **(c-d)** represent control and AD samples, respectively.

Fig. 5c-d shows the dynamic changes of circadian patterns of 8 representative genes from early to late stage of AD, including *PIGZ, PNPLA6, RAB11B,* and *CDT1* associated with STG, *SLC5A3,* and *ACAN* from PG, *OMA1* from IFG, and *SLC2A8* from FP. Interestingly, the amplitude of *PIGZ* is altered in the early stage and has a phase shift in the mid stage (Fig. 5c). *ACAN* has obvious rhythm at the early stage and a phase shift at the mid stage (Fig. 5d). *PNPLA6* and *RAB11B* shows attenuated amplitude in the early stage followed by loss of rhythm at the mid-stage. *CDT1* displays an obvious phase shift in the early stage. In addition, *SLC2A8* and *OMA1* gain the rhythmicity from early to mid-stage, while *SLC5A3* lost the rhythm starting from mid stage.

For better understanding the biological meaning of these 8 circadian genes identified above, we examined the genetic backgrounds of individual tissues and disease stages (Fig. 6). Since the single nucleotide polymorphisms (SNPs) are the major genetic information in Alzheimer’s disease studies, first, we started from identifying the stage-specific SNPs. We performed correlation study between the variant allele frequency of the known 42 AD-specific SNPs^68^ and expression of 8 genes that have circadian pattern change in their stages (Supplementary DataTable1). Then, we identified 22, 10, and 19 SNPs located near to 28, 10, and 25 genes. Among these, the SNPs located near to *ABCA7, CLU, CR1, EPHA1-AS1, MIR6843,* and *SORL1* were common across all AD stages. On the other hand, SNPs near to *ADAM10, ALPK2, BHMG1, CASS4,* and *PICALM* were early-stage specifically correlated with these 8 genes. These genes were enriched in the related pathways with ‘amyloid-beta formation’. During mid-stage, *CNTNAP2* was correlated. This gene is known as downregulated in the hippocampus of Alzheimer’s disease patients^69^. In the late-stage, *CNN2, MS4A6A, PILRA, SCIMP,* and *ZFP3* showed correlation with 8 genes. These genes were not enriched in any specific pathways. It seems the known AD-specific SNPs are related to our 8 circadian candidate genes in the early-stage to make worsen the condition of the patient. Next, to understand the biological impacts of these 8 genes, we performed correlation studies between the 8 genes and all expressed genes (1.0 ≤average TPM) per individual tissues and stages of MSBB cohort (Supplementary DataTable2). A total of 672 genes were significantly correlated with these 8 genes (Pearson correlation coefficient >0.9 and p-value < 1.0E-6). Out of 672 genes, 85 genes were matched with the tissue types and stages of the 8 genes, as shown in Fig. 6a. SLC2A8 has the largest number of correlated genes in FP tissue, including 61, 23, and 4 genes in the early, mid, and late stages, respectively.We performed gene set enrichment analysis focusing on 61, 23, and 4 genes in the early-, mid-, and late-stages using ToppGene^70^ as shown in Fig. 6b. Interestingly, those genes strongly correlated with *SLC2A8* in FP tissue and showed the anticipated enriched biological processes across different AD stages. For example, in the control group, 185 genes are involved in neuron differentiation and development with cell growth. However, in the early stage of AD, 61 correlated genes with *SLC2A8* are involved in the negative regulation of growth, which is opposite to the function in the control group. In the mid stage, 23 correlated genes has main function in the synaptic vesicle uncoating. This observation is consistent with the previous study about Clathrin-mediated endocytosis of synaptic vesicle in Alzheimer’s disease. The authors found that the synaptic activity-stimulated endocytosis enhanced the internalization of APP, resulting in the production and release of amyloid-beta (Aβ). In the late stage, the identified 4 genes has function in cardiovascular development. It has been reported that vascular dysfunction is likely to act synergistically with neurodegenerative changes to exacerbate the cognitive impairment in AD^71^. Further, a recent study indicated that older people with higher average blood pressure are also more likely to develop tangles and plaques in their brain than their peers ^72^. Furthermore, among 85 correlated genes with the matched tissue and stage with the changing circadian rhythms, 11 of them are AD-risk genes. Here, AD genes were downloaded from the International Genomics of Alzheimer’s Disease Project (IGAP^73^) and Accelerating Medicines Partnership-Alzheimer’s Disease (AMP-AD^39^). Those 11 genes include *AP2A1, AP3D1, CTIF, GAK, IL6ST, MINK1, PTPRS, RAB11FIP3, SLC2A8, SLC5A3,* and *TMEM184B.* Specifically, *MINK1* showed a significant correlation with *SLC2A8* in the FP tissue at the early and mid stages. Broce, *et al.* reported that *MINK1* is a gene with pleiotropic SNPs between AD and cardiovascular disease in a large ‘AD-by-proxy’ cohort from the UK Biobank and the expression of MINK1 was differentially altered in the postmortem AD brains^74^. In addition, we also revealed differential expression patterns of 6 of these 8 genes across different stages by one-way ANOVA. Specifically, *SLC5A3* and *RAB11B* showed differential expression patterns in PG and STG, respectively, consistent with their circadian rhythm changes in these tissues (Fig. 6c). Furthermore, we also analyzed the alternative splicing events and RNA A-to-I editing events in these 8 genes. However, there was no specific difference among AD stages. Based on our analyses, we infer that these circadian genes do not change randomly across pseudo time, but serve as intermediate genes in AD pathogenesis.

**Fig. 6.**
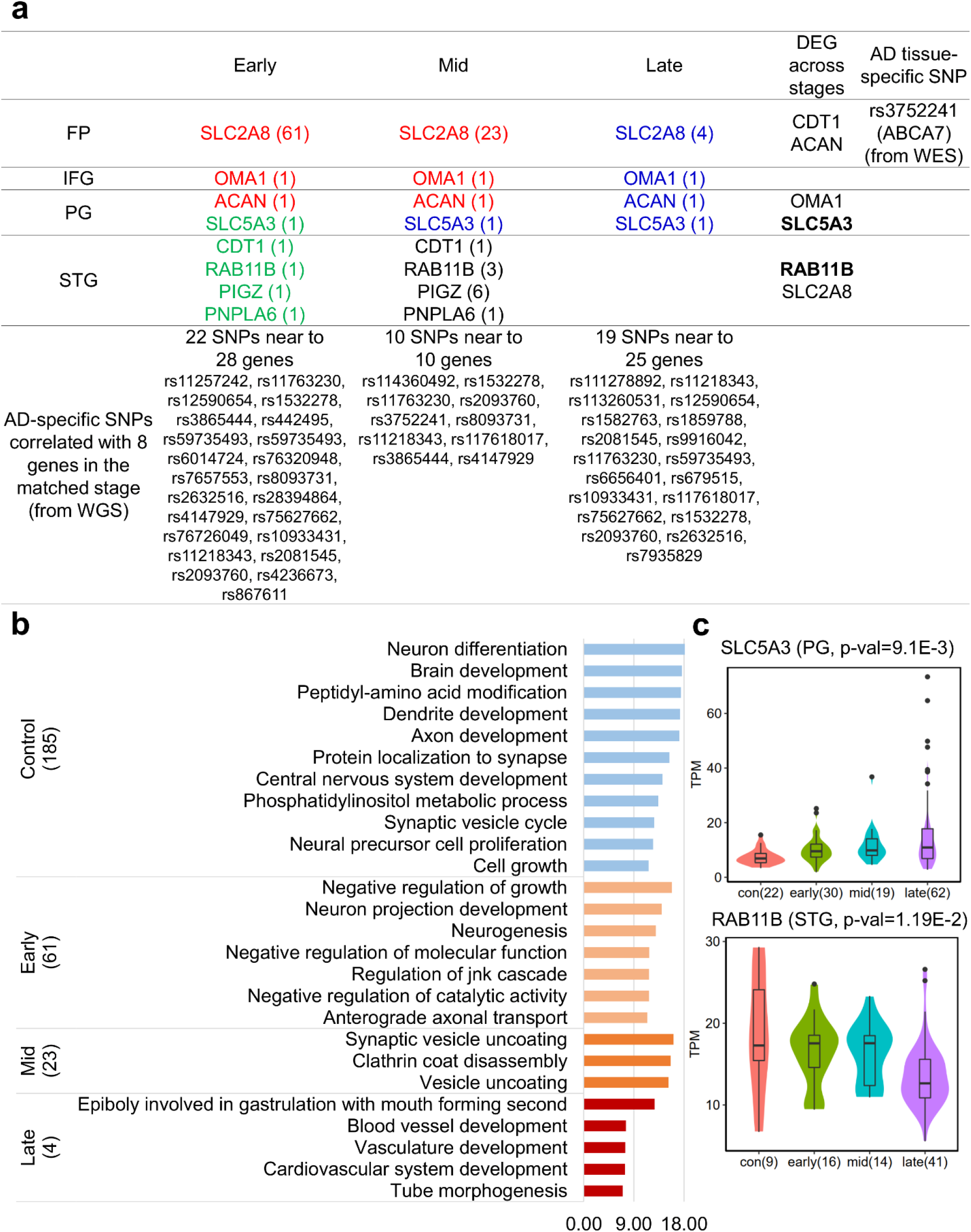
Genetic backgrounds of the 8 genes with circadian pattern changes in the MSBB cohort. **a,** Overall key genes among stages. The red fonts are the genes that gained the circadian rhythm compared to the previous stage. The blue fonts are the genes that lost the circadian rhythm. The green fonts are the genes that reduced the amplitude of circadian rhythm or had a phase shift. The bold fonts are the genes that showed the differential gene expression in the tissue matched with the circadian rhythm changes. SLC5A3 and RAB11B had differential gene expressions across stages. There were seven genes that have AD-specific SNPs associated with AD stages. **b,** Enriched biological functions of the correlated genes with SLC2A8 in FP tissue across stages. **c,** Expression difference across stages of SLC5A3 and RAB11B.

## Discussion

Recent findings have demonstrated that circadian dysfunction might be both a cause and an effect of neurodegenerative disorders during the AD progression^1,5,16,19,22,27,75^. Much less progress has been made in uncovering the molecular mechanism of how circadian rhythm links to AD development. The lack of time-course transcriptome data for AD has been a key barrier for translational medicine. Our ESOCVD system aims to address this deficiency by using global descriptors of gene expression and Bayesian approach to order unordered data. Based on the predicted circadian phase of each sample, we can order the untimed gene expression data and thus detect the circadian changes and their evolutions in the process of AD progression.

In this study, ESOCVD was applied to 20 untimed gene expression profiles in 16 brain regions of around 3000 AD patients from public transcriptome databases. Together, we examined up to 2391 genes (1691 seed genes and 700 PBCG genes) and observed 1) the distribution of the circadian rhythm changes in all the brain regions of late-onset AD; and 2) the evolution of the circadian rhythm changes in AD progression. Firstly, the preliminary test indicates that the outcomes of ESOCVD are reliable. We found that the inferred circadian patterns of 15 clock genes in the control group are consistent with the evidence reported in CGDB (Supplementary Table 2). Furthermore, we also found that the predicted circadian patterns of 43 PBCG genes were close to the evidences reported in CGDB (Supplementary Table 13). Secondly, our results revealed that loss or gain of rhythmicity occurred with higher frequency rather than the other three types of circadian variation patterns (Fig. 2 and Fig. 4). Thirdly, multiple well-known AD risk factors shows circadian alterations in at least 2 brain regions, e.g. *CR1, TREM2, PLD3,* and *HLA-DRB5, etc.* (Supplementary Table 9). Pathway-level analysis indicates that those circadian changes in AD are mainly associated with five brain regions, including SFG, STG, FC, TC, PC, *etc*. Analysis of circadian rhythm patterns of AD-related clock control genes in all brain regions revealed that 6 of them lost rhythm in certain regions (Fig. 3). Gene expression analysis of AD mouse brain tissues further substantiated the predictions of the ESOCVD model, which provides actionable knowledge for clinical treatment. Thirdly, we developed a systems biology approach to infer GRN to reveal the associations between AD development and circadian dysfunction. Our model fully utilizes the upstream and downstream regulatory information around TF and integrates the functional linkage data and gene expression data. By systematically optimizing the pathway network topology with functional linkage information and gene expression data, our model revealed three TF networks, demonstrating how the upstream of circadian rhythm TF modulate downstream target genes and thus promote AD progression (Fig. 4c-e). We analyzed the evolution of circadian rhythm changes in AD development. Through the integreation of other omics data, eight representative circadian genes were selected to explore their genetic background and dynamic changes across different stages. Our analysis shows that those 8 circadian genes may serve as intermediators in AD pathogenesis

ESOCVD modeling is initialized at a random starting point and converges to the global optimal solution. In order to improve the searching efficiency of our model, a novel heuristic search strategy was designed and incorporated into ESOCVD (Supplementary Fig. 9). Supplementary Fig. 10 shows that the likelihood calculation of a new sequence *S_ML_* generated from the initial sequence *S_bg_* by exchanging the positions of two events only need to consider a few top-ranked cases of *S_bg_* (Fig. 1). It indicates that the knowledge gained in the current step can be used to guide the new search. To against local minima, the optimal sequence for each dataset was obtained after multiple runs. To ensure good convergence, the ESOCVD model was repeated 50, 100, and 200 times for each profile staring from different initial points. Supplementary Fig. 11 indicates that our model displays good convergence. 200-runs is enough to locate the global optimal solution (Supplementary Table 14).

Individual differences between patients may make it difficult to compare baselines and periodic patterns of gene expression. For each dataset, we normalized gene expression profiles within and between samples before model training. In addition, we used 10-fold cross-validation to evaluate the impact of individual variation on the consistency of the optimal solution (see the details in “**Methods**”). Supplementary Fig. 12 shows that the similarity between the optimal sequence fitted to each fold and the sequence fitted to the whole data set is high.

ESOCVD outperforms other typical approaches. As a unique bioinformatics tool for oscillating rhythm detection on untimed gene expression data, CYCLOPS^38^ shows better efficiency and robustness than other tools, e.g. OSCOPE^37^, and ZeitZeiger^34^, *etc*. In this study, we mainly compared our ESOCVD with CYCLOPS on timed and untimed transcriptome datasets. Firstly, we applied both methods on three time-course transcriptome datasets (GSE39445, GSE48113, and GSE56931) and compared the accuracy on the prediction of the circadian phase. Our comparison result indicates that the prediction accuracy of ESOCVD is better than CYCLOPS (Supplementary Fig. 13). Secondly, we applied both methods on four untimed AD-related datasets with small to large scale to test the reproducibility of the estimated optimal sequence of events. The 10-fold cross-validation revealed that ESOCVD had the better reproducibility of the estimated optimal sequence of events (Supplementary Fig. 12). Furthermore, Supplementary Fig. 14 presents two typical cases of loss rhythmicity detected by CYCLOPS and ESOCVD, respectively. The performance of ESOCVD was close to CYCLOPS for the large size datasets (Supplementary Fig. 14a). Supplementary Fig. 14b shows that CYCLOPS trends to obtain the circadian phases around 12h and misses at least a half cycle. Moreover, our analysis revealed that the number of samples greatly limited the performance of CYCLOPS. The performance is significantly reduced if the sample size is relatively small (Supplementary Fig. 15), so that the estimated circadian phases of samples cannot fill in underrepresented time of day^38^, which leads to the circadian patterns in a single cycle cannot be completely detected. The above findings indicate that CYCLOPS is more sensitive to the sample size rather than ESOCVD.

There are several limitations of our ESOCVD model. Light-dark cycle and sleep patterns of patients from which the RNA data may influence gene expression and thus obscure waveform and amplitude of temporal fluctuations. Our model searched for the optimal periodic rhythm by fitting the gene expression across all subjects, therefore, it represents the overall trend of circadian rhythm in a patient population. The selection of seed genes may bring some bias on model outcomes. To deeply understand the association between circadian dysfunction and AD progression, we will further identify the seed genes suitable for ESOCVD modeling based on reported data and knowledge. Using 12 eigengenes (global descriptors) and 24 events, the resolution of the predicted circadian phase is 1 hour. Increasing both the number of eigengenes and the events can assign the samples more evenly distributed within the entire cycle. Given the relative positions of two events for a certain eigengene in an event sequence, the current model needs to consider two possible periodic patterns (Fig. 1), which will largely increase the computational cost in the likelihood calculation.

## Methods

### Identification of “seed genes”

Firstly, we identified 8504 circadian genes that are more likely to cycle in multiple tissues by using the data in literature^41^ via JTK-cycle^35^. Secondly, we further screened out 1670 brain-related circadian genes, including 15 well-known core clock genes (CLOCK, PER1, PER2, *etc*.). Finally, we combined above 1670 genes with 21 AD-related clock-controlled genes^45–47,50,51^, and thus obtained 1691 seed genes, which are considered as the pacemakers in the clock system of AD. Considering the fact that the expression profiles of the seed genes are still high-dimensional, we use SVD to reduce the dimensionality of the data and extract “eigengenes”, characteristic circadian expression patterns, which span the global expression profiles^38,76,77^. We normalized each patient sample on SVD matrix by subtracting the mean of control group and dividing by the standard deviation of control group. Finally, our controls have a mean of 0 and a standard deviation of 1, the patients won’t have a mean of 0 or standard deviation of 1, because they are z-scored relative to the controls and they should have some disease signal. Besides the 1691 seed genes, we also explore 857 PBCG genes and their circadian changes (Supplementary Fig. 1).

### Five patterns of circadian variation

In this study, we consider five types of circadian alterations between control and AD groups: (1) amplitude change (AC) in AD: a decrease but not the complete loss in amplitude (2-fold reduction); (2) a significant loss of rhythmicity (LR) in AD; (3) a gain of rhythmicity (GR) in AD but no significant rhythm in control; (4) base shift (BS): a change in levels of expression but not necessarily a change in rhythm; (5) phase shift (PS): no significant changes on the amplitude and mean level but shift the phase at least ¼ cycle (6 hours). Based on the above five modes, we can identify the clock genes and the output genes with circadian alteration in AD, respectively.

### ESOCVD modeling

We formulate the modeling underlying ESOCVD as groups of samples with distinct circadian patterns of eigengene evolution. The eigengene evolution is described as a periodic z-score model in which each eigengene follows an oscillating trajectory across a common timeframe (24 hours). Our model is based on the concept ‘event’ in the event-based models used for disease progression^78,79^. Different from a unique z-score value^78,79^ or linear z-score series^80^ described the monotonic transition from a normal to an abnormal level, ESOCVD reformulates the events of each eigengene as continuous periodic dynamics. The model fitting is implemented with a Bayesian approach to search the optimal ordering of the events (sequence **S**) for estimating the possible circadian phase of each sample.

### Mathematical basis of ESOCVD

Traditional event-based models describe disease progression as a series of events, where each event corresponds to a biomarker transitioning from normal to abnormal level^78,81^.

However, the instantaneous normal to abnormal events cannot depict the dynamics of the biomarker evolution trajectory. Young, *et al.* use events that represent the linear accumulation of a biomarker from one z-score to another, to uncover data-driven disease phenotypes with distinct temporal progression patterns^80^. For biomarker *i*, a set of events was represented as a set of monotonically increasing z-scores *z*_*i*1_, *z*_*i*2_, *z_iR_* indicating that the samples with larger expressions would be expected as severe. The limitation of the above methods is that they cannot address the non-monotonic biomarkers. Our developed ESOCVD uses the periodic z-score strategy (Fig. 1). We assume that the model works on 12 circadian eigengenes, which was extracted from the original expression matrix of seed genes with SVD^76,77^. For the eigengene *i* (*i* = 1, we define two events *E*_*i*1_ and *E*_*i*2_, with z-score *z*_*i*1_ and *z*_*i*2_, which represent the ‘crests’ or ‘troughs’ in a single cycle. There are two cases (evolution patterns) need to be considered only given the relative position of *E*_*i*1_ and *E*_*i*2_ (Fig. 1). Totally, we have *N* = 2 * *I* = 24 events locating the *x* axis 1-24 (Fig. 1). The ‘distance’ between any pair of *z*_*i*1_ and *z*_*i*2_ is half cycle (12h). Therefore, a candidate solution of ESOCVD, event sequence **S**, needs to follow the constraints shown as below (**Eq. (1–3)**):

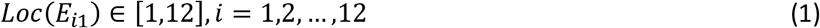

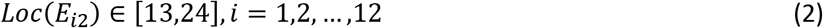

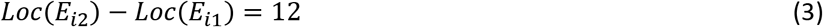

where function *Loc()* defines the position of a z-score event in the sequence **S. Eq. (1–3)** assume that the event *E*_*i*2_ occurred later than event *E*_*i*1_. The occurrence of an event *E_i_*, is informed by the measurements *x_i,j_* of eigengene *i* in subject (sample) *j, j* = 1 …*J*. The dataset *X* = {*x_i,j_*|, … = 1, …, *I,j* = 1,…,|} is the set of observations of each eigengene in each subject. The aim of ESOCVD is to find the most likely ordering (sequence **S** in Fig. 1) of the events that maximize the data likelihood *P*(*X|S*). We consider a single cycle as a continuous time axis, *t*, which we choose to go from *t* = 0 to *t* = 1. At each stage *k*, which goes from *t* = *k*/(*N* + 1) to *t* = (*k* + 1)/(*N* + 1), and a z-score event *E*_*i*1_ (*q* equals to 1 or 2) occurs. Given the positions of *E*_*iq*_ and *E*_*i*2_ in sequence **S**, the eigengene *i* at the beginning of stage *k* can be determined by **Eq. (4)**:

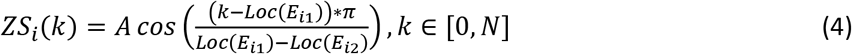

The model likelihood *P*(*X|S*) can be represented as **Eq. (5)**:

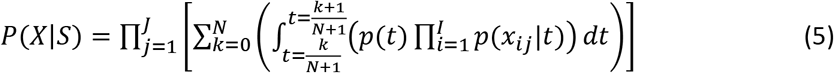

where *p*(*x_ij_*|*t*) = *NormFDF*(*x_ij_; g_i_*(*t*); *σ_i_*). *NormFDF*(*x,μ,σ*) is the normal probability density function, with mean *μ* and standard deviation *σ*, evaluated at *x. p*(*t*) follows a uniform distribution, indicating that a prior individual is equally likely to belong to any stage along the oscillating pattern. *g_i_*(*t*) is a line segment of the form *g_i_*(*t*) = *a_i_t* + *b_i_*. It means the change of *g_i_*(*t*) is linear if 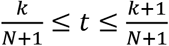. Thus, we have *g_i_*(*t*) = *ZS_i_*(*k*) if *k = t* * (*N* + 1).

### Model fitting

Supplementary Fig. 9 presents the flowchart of model fitting in our ESOCVD. Model fitting requires simultaneously the stage membership, and the posterior distributions. The sequence **S** is initialized randomly. And then, this sequence is optimized by using our hybrid search algorithm. The hybrid search strategy combines global search and local search to find the optimal position of each event in the sequence. In our simulation, the sequence **S** is optimized from 12 different random starting sequences to find the maximum likelihood solution.

### Hybrid Search strategy

The hybrid search algorithm involved in ESOCVD model starts from an initial sequence *S_q_*, then performs global search and local search, and finally return an optimal sequence *S_ML_* (Supplementary Fig. 10). For simplicity, we used the integer 1-24 to represent 24 original events: *E*_1,1_,*E*_2,1_, …,*E*_12,1_,*E*_1,2_,*E*_2,2_, …, *E*_12,2_, which are associated with 12 eigengenes (Fig. 1). The inferred sequence *S_ML_* (the same as the index sequence in Fig. 1) obtained from our model represents the index of each event in the optimal sequence S relative to the original event profiles. Constrained by **Eq. (1–3)**, each valid sequence *S_p_* satisfies two conditions: (1) the sub-sequence *S_q_*(1), …,*S_q_*(12) is a permutation of the integers from 1 to 12; (2) the sub-sequence *S_q_*(13),…,*S_q_*(24) was determined *S_q_*(1), …, Eq (12) via **Eq. (3)**. The calculation procedure includes the following steps:

**Step 1**: Input an initial sequence *S_q_*. To guarantee the variety of start points, we set 1 ≤ *q* ≤ 12. It indicates the first event in sequence *S_q_* is event *E*_*q*1_.
**Step 2**: Global search is performed based on sequence *S_q_*. A new sequence *S_g_* can be generated through three step3: 1) *S_g_*(1) = *S_p_*(1); 2) permute the elements *S_q_*(2), …, *S_q_*(12) of *S_q_* and assign them to *S_g_*(2), …,*S_g_*(12); (3) calculate *S_g_*(13),…,*S_g_*(24) with **Eq. (3)**. Then, calculate the likelihood of each *S_g_* on data *X*, and screen the sequence *S_bg_* with maximal likelihood. In our simulation, we totally generate 100 new sequence *S_g_* from an initial sequence *S_q_*.
**Step 3**: By using a heuristic search strategy, local search is further performed based on the results of the global search (Supplementary Fig. 9-10). In this step, we exchange the positions of any two events (*E*_*i*1_ ↔ *E*_*j*1_ and *E*_*i*2_ ↔ *E*_*j*2_) in sequence *S_bg_* each iteration and generate a new sequence *S_L_*, and then evaluate if the likelihood of *S_L_* is larger than *S_bg_*. Given the relative positions of two events of each eigengene *i* (*E*_*i*1_, and *E*_*i*2_) in a sequence S, there are two possible circadian patterns that need to be tested. Totally, we need to test the likelihood values in all the 2^12^ = 4096 possible cases to determine the maximal likelihood for a sequence **S**, which may lead to high computational cost. Therefore, we proposed an approximate search strategy to speed up the calculation of likelihood. The *rationale* is: if the distance of two events (*E_i_* and *E_j_*) in sequence *S_bg_* is very close, the new sequence *S_L_*, generated from *S_bg_*, by exchanging *E_i_* and *E_j_*, will overlap the top cases of *S_bg_* with a big ratio. Top cases are a subset of all the 4096 cases, which will return good likelihood values. To prove the effectiveness of this rational, we selected GSE125583 and MSBB datasets to test. Firstly, we ranked all the 4096 cases of sequence *S_bg_* according to the descending order of 4096 likelihood values. Secondly, we ranked all the 4096 cases of a new sequence *S_L_* in the same way. Thirdly, we calculated the overlap of cases with top 100 if the distance between *E*_*i*1_ and *E*_*j*1_ is 1. Similarly, calculate the overlap of cases with top 200 if the distance between between *E*_*i*1_ and *E*_*j*1_ is 2. Our result shows that around 90% of the top 100 cases of *S_bg_* also rank within top 100 for *S_L_*, which indicates that we can only use the top 100 cases of *S_bg_* to calculate the likelihood of *S_L_* if the distance between *E*_*i*1_ and *E*_*j*1_ is 1 (Supplementary Fig. 16). In the meantime, the overlap ratio will be decreased if the distance between *E*_*i*1_ and *E*_*j*1_ is large. Therefore, our approximate strategy sharply reduces the computational cost of likelihood calculation. In our model, we use a 12-bit binary number to represent a possible case for the circadian patterns of 12 eigengenes. For example, 101000000000 means only the event *E*_1_ and *E*_3_ in a sequence S follow the first case of their circadian patterns, and the other events follow the second case.
**Step 4**: screen the optimal sequence *S_ML_*. After performing the local search, we compare all the sequences *S_L_* with *S_bg_*, and determine the optimal sequence *S_ML_* with the maximal likelihood.

### Convergence

As described in the above section, the model fitting performs a search optimization from different starting points and chooses the sequence with maximal likelihood as the final solution. In this study, we repeat the ESOCVD model 50, 100, and 200 times for each dataset in parallel on Texas Advance Computing Center (TACC) to against the local minimum of model fitting. We find that ESOCVD shows good convergence: runs from different start points typically converge to a solution.

### Cross-validation

To evaluate the consistency of the inferred optimal sequence, we performed 10-fold cross-validation by dividing the data into ten folds and re-fitting the model to each subset of the data, with one of the folds retained for testing each time. And then, we compute the similarity of any two optimal solutions inferred from the model fitted to each subset and the model fitted to the whole dataset, respectively.

### Reconstruction of the GRN network

To reveal the association between circadian dysfunction and AD development, we developed a systems biology approach to reconstruct the circadian rhythm TF regulatory network (Supplementary Fig. 3). The calculation includes the following steps:

**Step 1**: *Inference of pairwise transcriptional regulations*. Based on the AD risk factors were detected as circadian alterations, we searched all validated TF-TG pairs from the public platform TRRUST^61^. There are two situations: 1) If the risk factor with circadian variation is a TG, we need to firstly screen out all the possible TFs for this TG and only select the TFs with the same type of circadian changes as TG; 2) If the risk factor is a TF, we selected all the possible TGs from TRRUST.
**Step 2**: *Inference of modulatory triplet*. To obtain the modulator (cofactor)-TF-TG triple regulatory relationships, we firstly identify all the possible modulators of the TF-TG pairs. In this study, we restrict the candidate modulators in the cofactor gene set that interact with TF in functional linkage network from STRING database^66^. We mainly focus on the top-ranked candidate cofactors, which were reported in literature and were validated by other researchers in the previous experiments.
**Step 3**: *Integrating upstream and downstream regulations*. The upstream and downstream regulations for all the TFs in the same tissue need to be integrated as a generic pathway map^67,82^. Please see two examples shown in Supplementary Fig. 4.
**Step 4**: *Inference of circadian rhythm TF-specific pathways by TILP.* To infer TF-specific pathway networks, the well-established TILP^67^ approach is used to modeling the generic pathway map with gene expression profile. The constraint system involved in TILP is designed to represent the relationship between the states of two linked nodes and their edge. Nodes in the network represent proteins/genes with their discrete values: −1, 1, or 0, where −1, 1, and 0 stands for “down-regulation”, “up-regulation”, and “no-change”, respectively. The signaling events (connections) are encoded by the operations on the nodes, which also have discrete values (‘promotion’ (1) or ‘inhibition’ (−1)). By fitting the TILP model of a generic pathway to the states of observed nodes via the constraints, we can remove the inconsistent connections and obtain TF-specific pathways. In this study, we used the fold change (FC) of the mean value of a gene in AD to control to get the discrete state of each node: if FC>1.2, “up-regulation”; if FC<1/1.2, “down-regulation”; otherwise, “no-change”.

### Real-time qPCR

APPswe/PSEN1dE9 (APP/PS1) mice (JAX: 034829) mice were maintained and treated according to IACUC guidelines, and the procedures were conducted as described in animal protocols approved by the IACUC of the University of Texas Health Science Center at Houston (UTHSC-H). APP/PS1 and wild-type littermate mice were collected at the age of 19-20 months at two circadian time points (ZT6 and ZT18). RT-qPCR analysis was conducted as previously described ^83,84^. Briefly, cortex tissues were isolated from the whole brain and pulverized. Total RNA was extracted from frozen cortex powder by using TRizol (Invitrogen). One μg of extracted RNA was used for cDNA synthesis. Gene expression was analyzed by using QuantStudio 7 Flex Real-Time PCR. Primers for qPCRs are: Atxn1, forward: CACGGTCATTCAGACCACACA, and reverse: GGTAGCCGATGACAGGAGGTT; Atxn3, forward: TGTCTTGTTACAGAAAGATCAG, and reverse: GTTACAAGAACAGAGCTGACT; Scna, forward: TGACAGCAGTCGCTCAGA, and reverse: CATGTCTTCCAGGATTCCTTC; Scnb, forward: GGAGGAGCTGTGTTCTCTGG, and reverse: TCCTCTGGCTTCAGGTCTGT; Bace2, forward: TGAGGACCTTGTCACCATCCCAAA, and reverse: TGGCCAAAGCAGCATAAGCAAGTC; ApoE, forward: AACCGCTTCTGGGATTACCT, and reverse: TGTGTGACTTGGGAGCTCTG; Mapk1, forward: GGAGCAGTATTATGACCCAAGTGA, and reverse: TCGTCCACTCCATGTCAAACT; Fos, forward: CAGAGCGGGAATGGTGAAGA, and reverse: CTGTCTCCGCTTGGAGTGTA; Clu, forward: CAGTTCCCAGACGTGGATTT, reverse: TCATCTTCAGGCATCCTGTG.

### Code availability

All the raw data used in this study are available from GEO and AMP-AD. Source code is available at github: https://github.com/JakeJiUThealth/ESOCVD_1.0.

## Acknowledgments

This work was supported by National Institutes of Health U01CA166886, U01AR069395-01A1, R01CA241930 and R01GM123037 (Zhou). Welch Foundation (AU-1971-20180324) and NIH/NIGMS (R01GM114424) to S.-H.Y., The Welch Foundation (AU-1731-20190330) and NIH/NIA (RF1AG061901, R56AG063746 and R01AG065984) to Z.C. Funding for open access charge: Dr Carl V. Vartian Chair Professorship Funds to Dr. Zhou from the University of Texas Health Science Center at Houston. The funders had no role in study design, data collection and analysis, decision to publish, or preparation of the manuscript.

## Author contributions

Z.J. and X.Z. conceived and designed the computational framework. Z.J. implemented programming code and wrote the manuscript. D.S., P.K., S.W, M.Y analyzed the data. S.Y., Z.C. designed the gene expression analysis which was performed by E.K. and M.W. W.Z. advised on the description of some analysis. All authors contributed to reviewing and editing of the report.

## Additional information

**Supplementary Files** accompanies this paper at the following links:

Supplementary File1: https://ccsm.uth.edu/ESOCVD/SupplFile1.rar.

Supplementary File2: https://ccsm.uth.edu/ESOCVD/SupplFile2.rar.

Supplementary File3: https://ccsm.uth.edu/ESOCVD/SupplFile3.rar.

Supplementary DataTable1: https://ccsm.uth.edu/ESOCVD/SupplDTb1.xlsx.

Supplementary DataTable2: https://ccsm.uth.edu/ESOCVD/SupplDTb2.xlsx.

