## Supplemental Tables, Figures for "A novel bioinformatics approach to reveal the role of circadian oscillations in AD development"

### Supplementary Figures

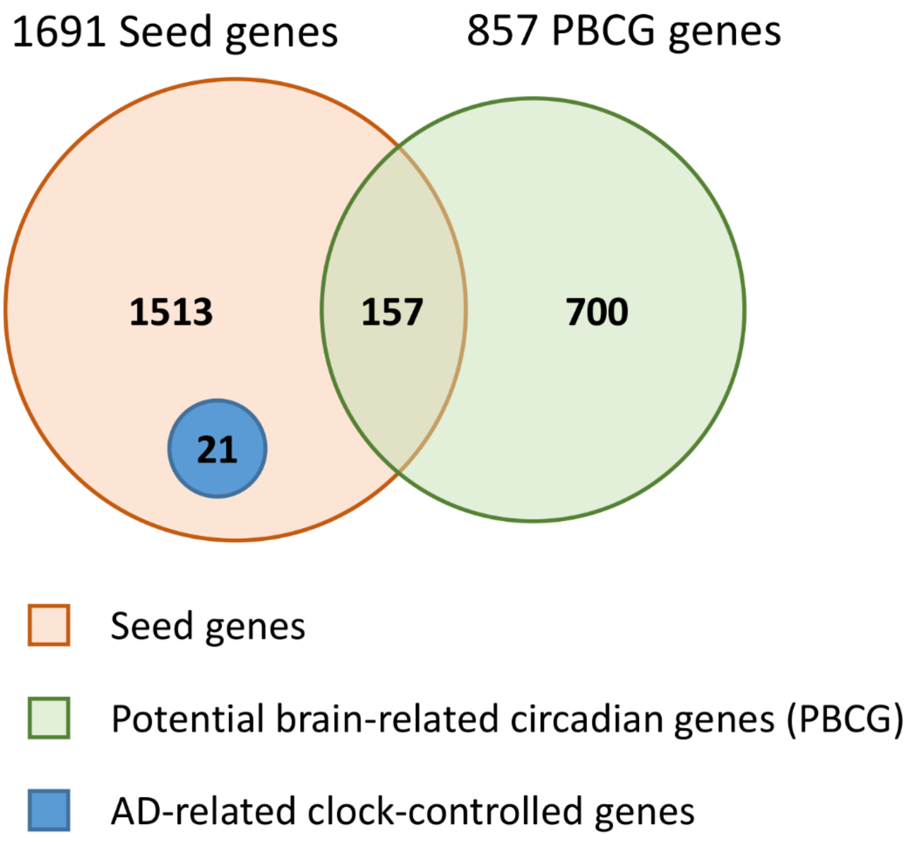


**Supplementary Fig. 1** The seed genes and PBCG genes studied in this work.

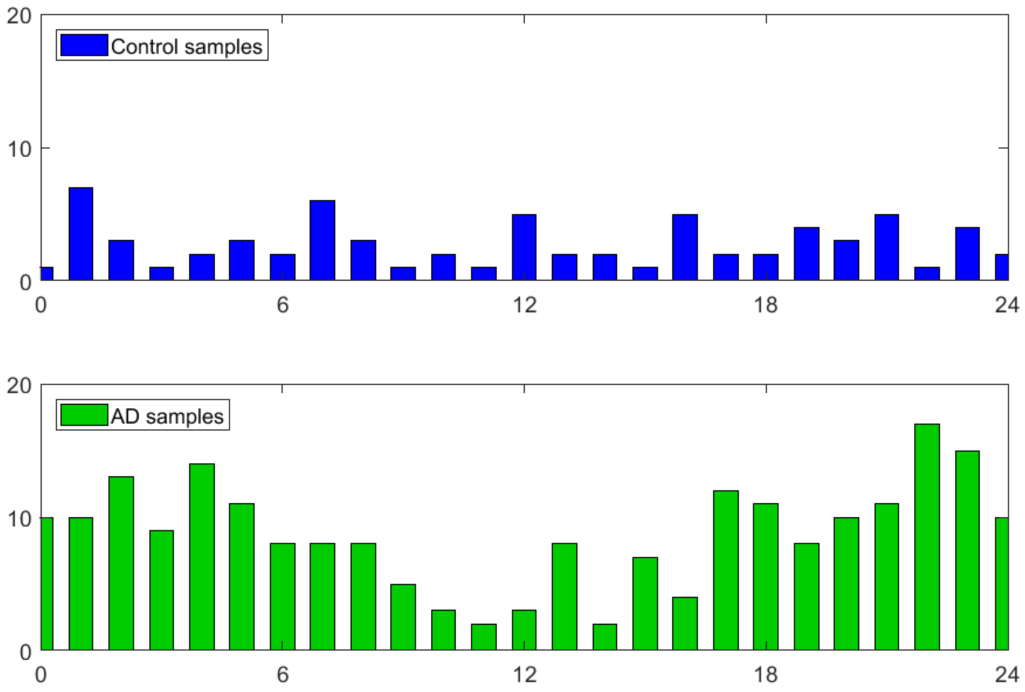


**Supplementary Fig. 2**. The prediction of subjects in GSE125583 belong to each of the circadian phase.

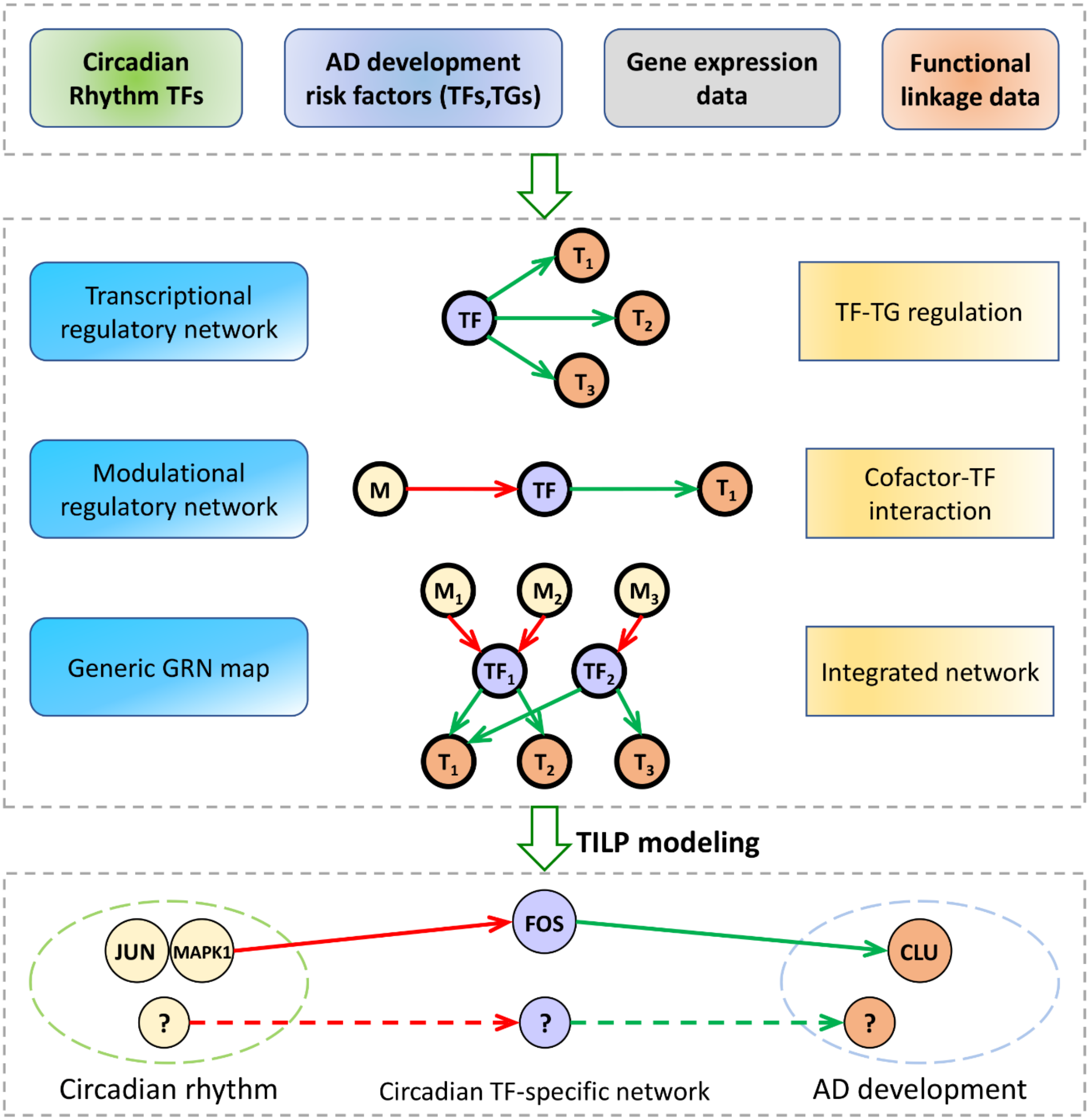


**Supplementary Fig. 3**. The network construction workflow for AD development and circadian rhythm process.

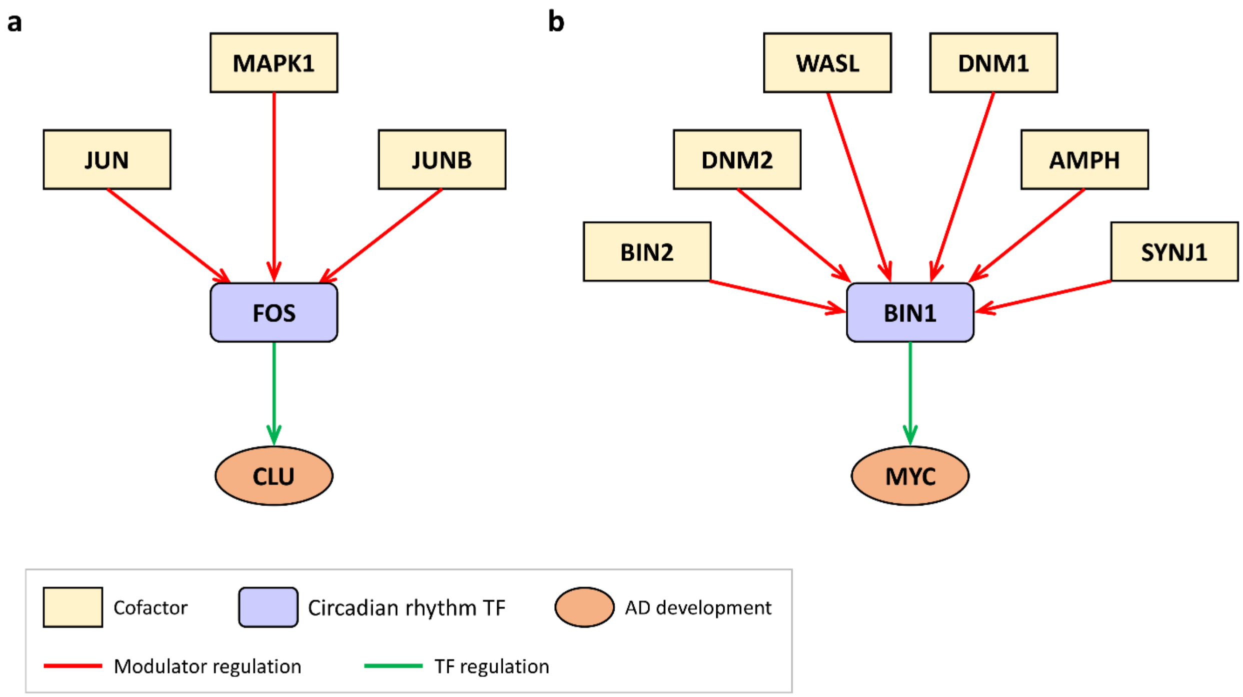


**Supplementary Fig. 4** Two generic pathway maps associated with AD risk factor (**a**) CLU and (**b**) BIN1, respectively.

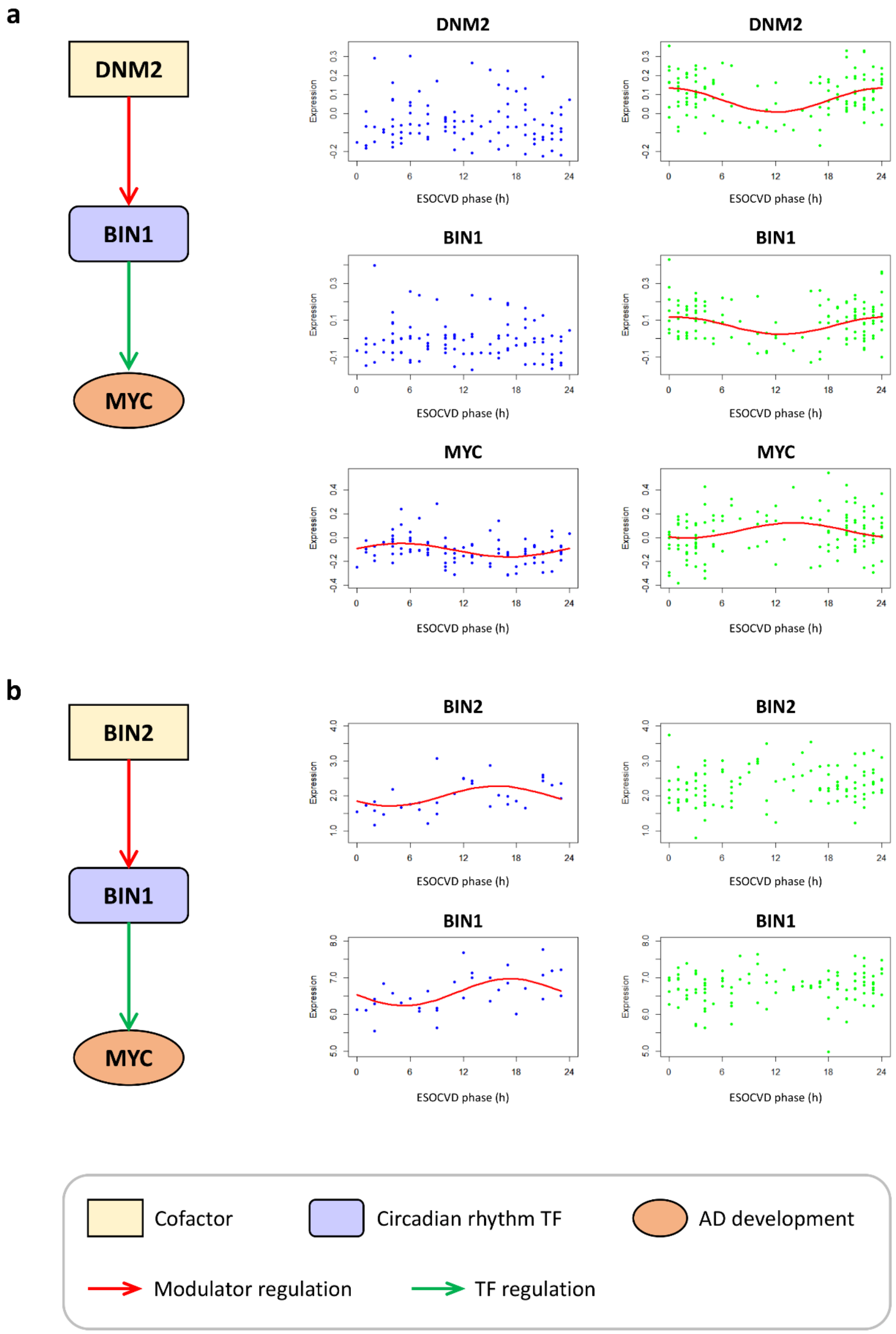


**Supplementary Fig. 5** Inferred BIN1 regulatory networks in VC and STG.

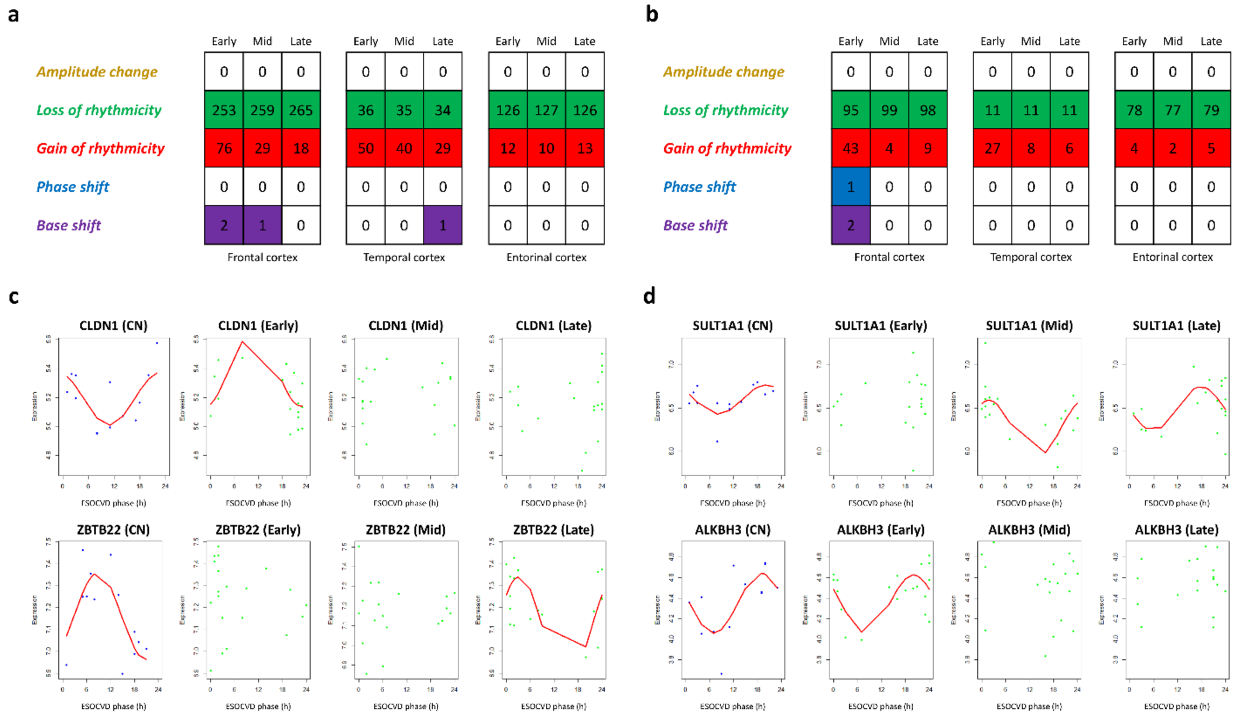


**Supplementary Fig. 6** ESOCVD analysis of circadian dynamics in three regions during AD progression. Expression data from GSE131617 was used. **a**, the number of seed genes with circadian changes in three brain regions detected by ESOCVD. **b**, the number of PBCG genes with circadian changes in three brain regions detected by ESOCVD. **c-d**, expression of four selected seed and PBCG genes as a function of ESOCVD phase. CLDN1 and SULT1A1 are detected from FC. ZBTB22 and ALKBH3 are associated with TC and EC, respectively.

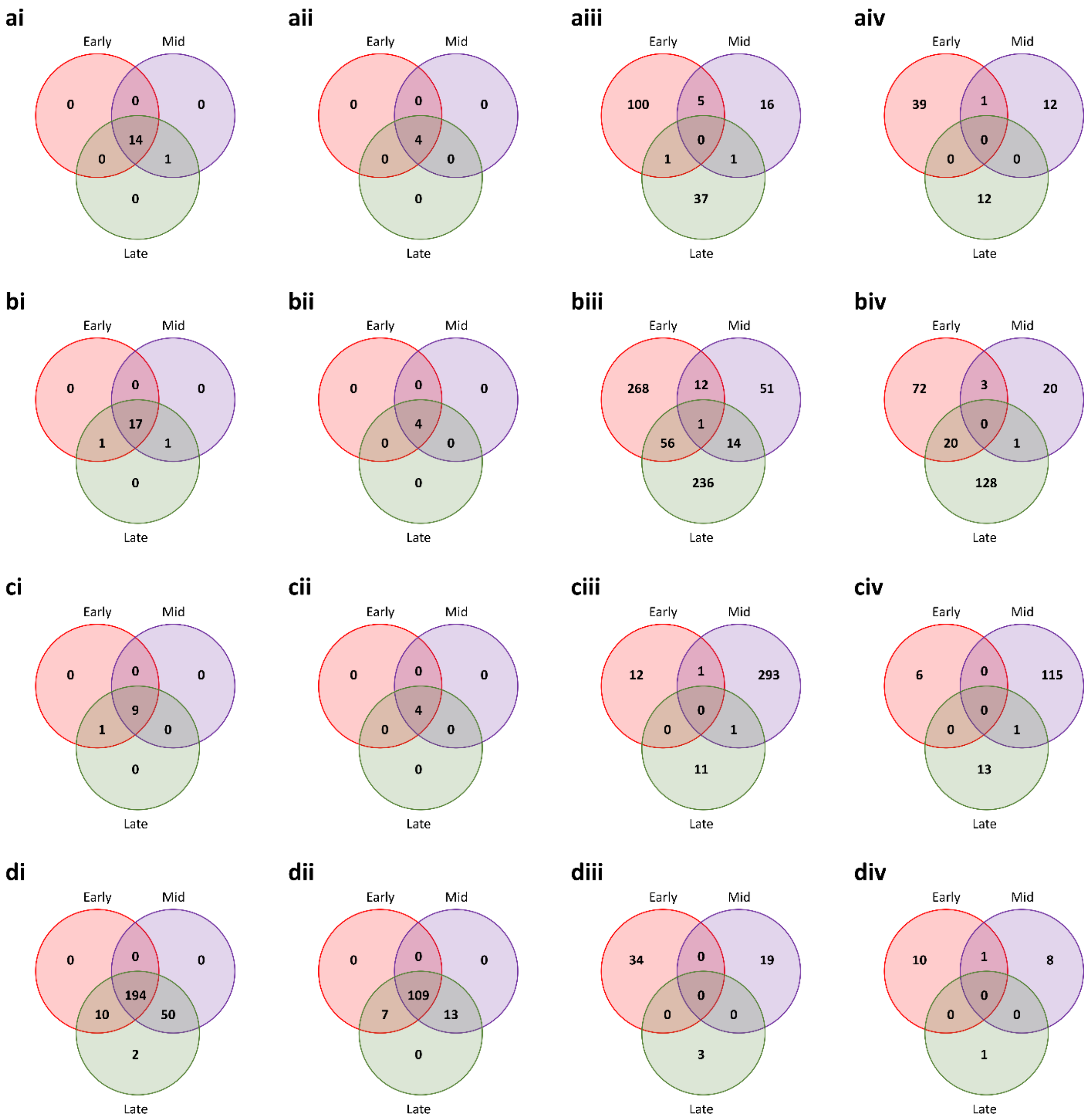


**Supplementary Fig. 7** Overlap between genes in early, mid, and late stage in region (**a**) FP, (**b**) PG, (**c**) IFG, and (**d**) STG. The seed genes (i and iii) and PBCG genes (ii and iv) with loss (i and ii) and gain (iii and iv) of rhythmicity are presented. Expression data from MSBB is used.

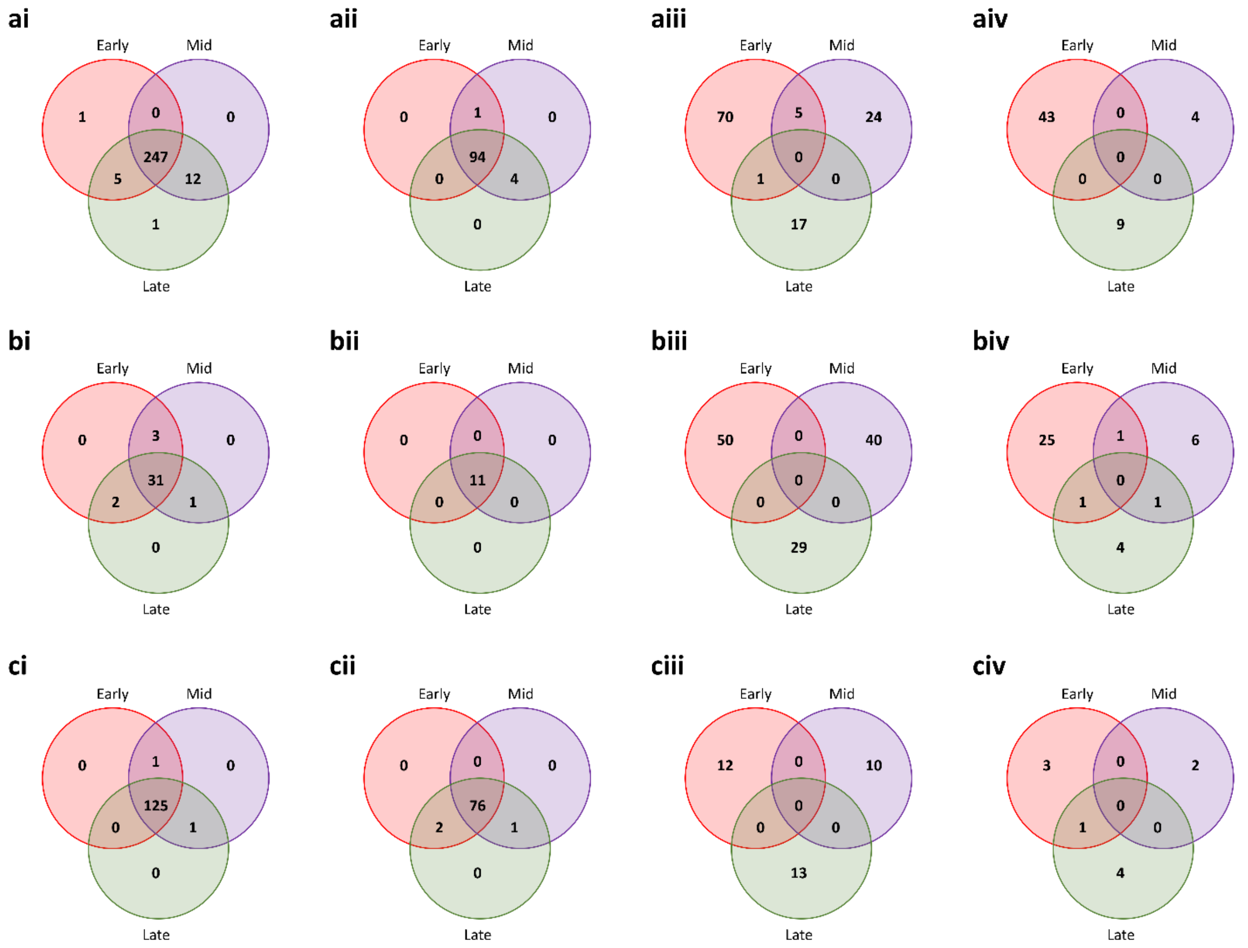


**Supplementary Fig. 8** Overlap between genes in early, mid, and late stage in region (**a**) FC, (**b**) TC, and (**c**) EC. The seed genes (i and iii) and PBCG genes (ii and iv) with loss (i and ii) and gain (iii and iv) of rhythmicity are presented. Expression data from GSE131617 is used.

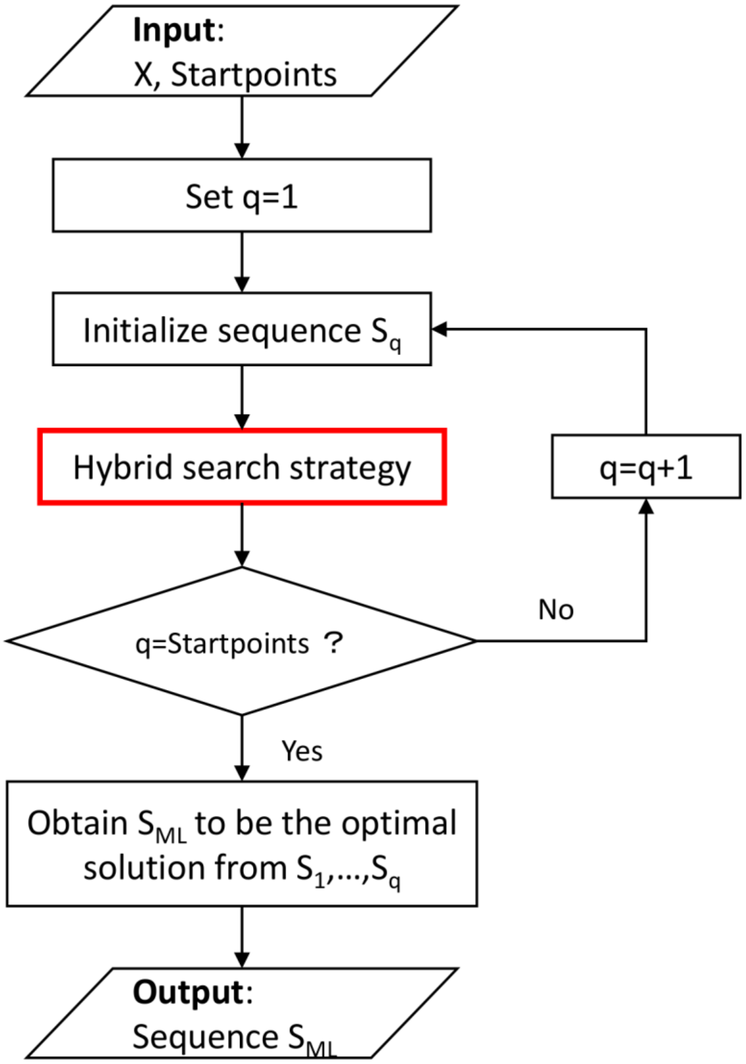


**Supplementary Fig. 9** Flowchart of the processes involved in the ESOCVD model fitting.

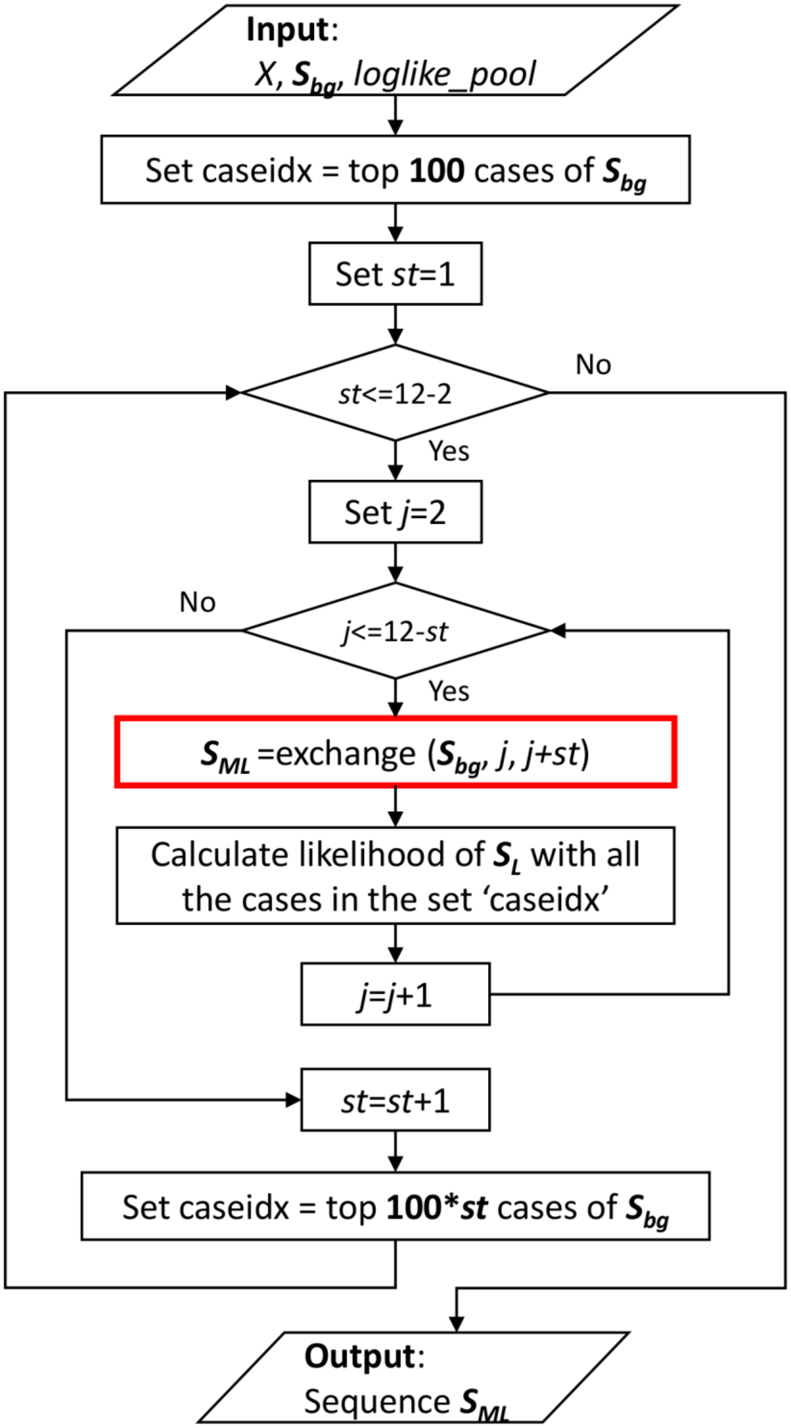


**Supplementary Fig. 10** Flowchart of the hybrid search strategy involved in ESOCVD model.

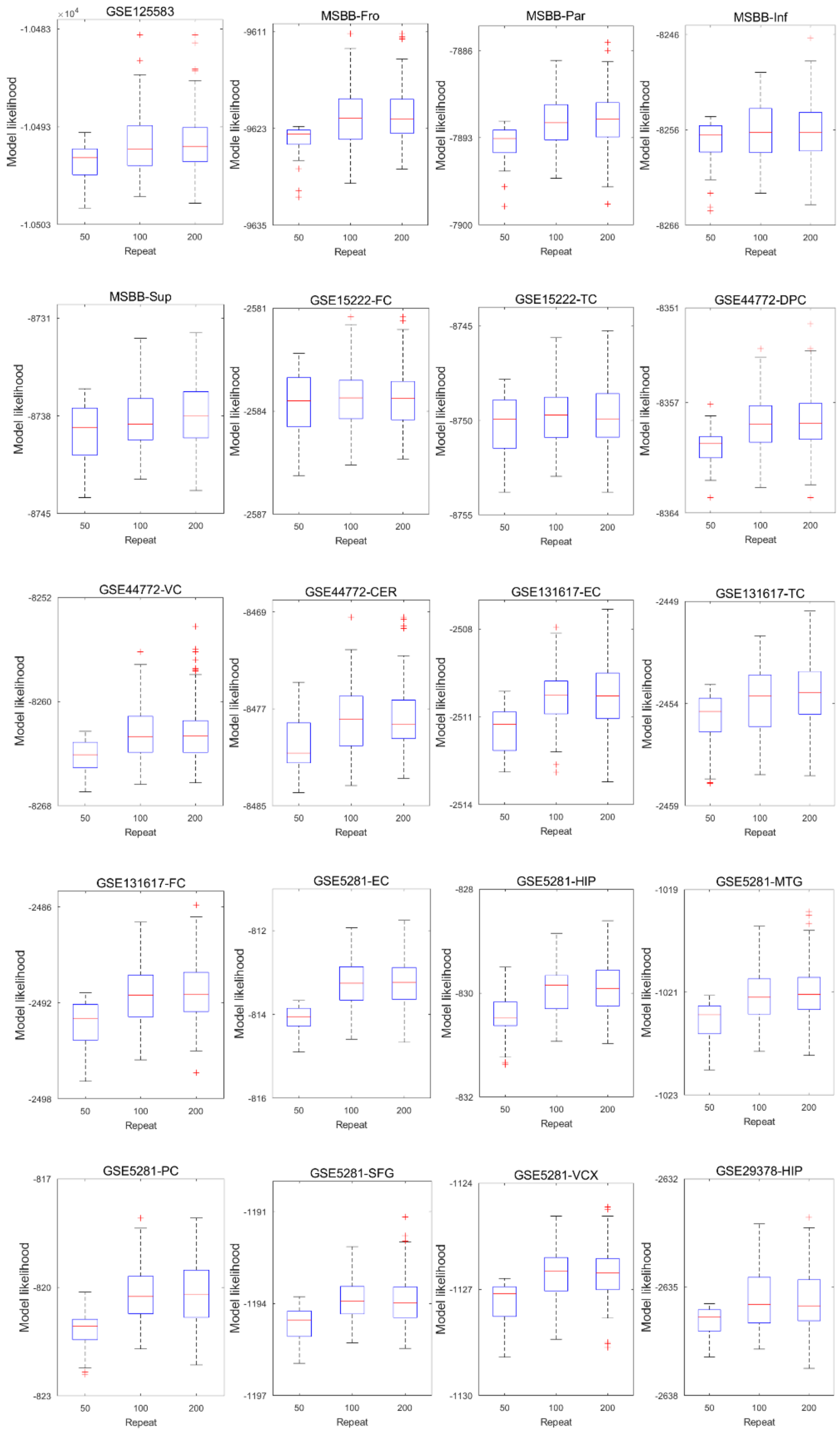


**Supplementary Fig. 11** The distribution of maximal likelihood values by running ESOCVD 50, 100, and 200 times.

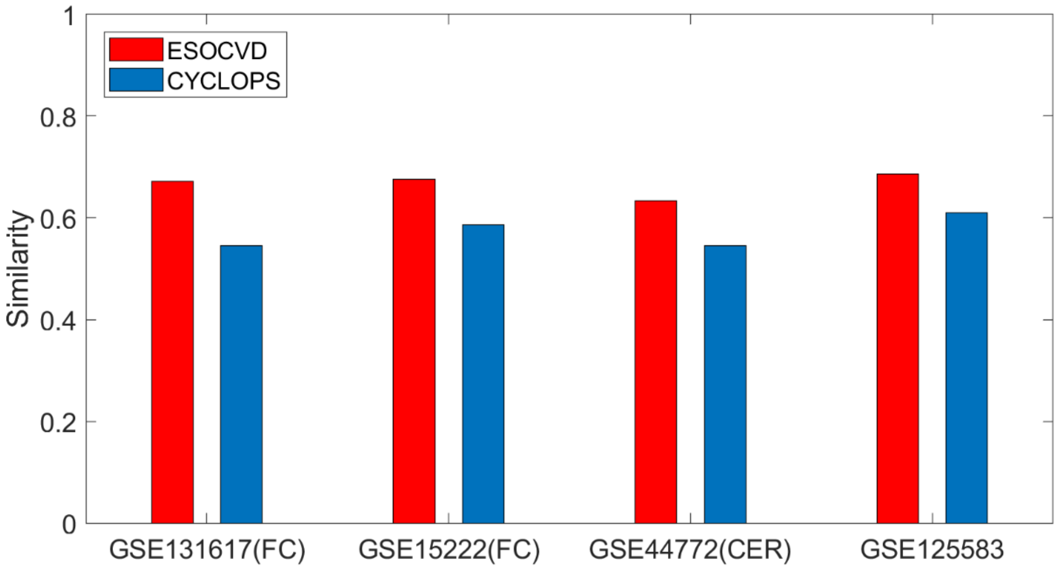


**Supplementary Fig. 12** The comparison between ESOCVD and CYCLOPS on four untimed transcriptome datasets of AD from small to large scale.

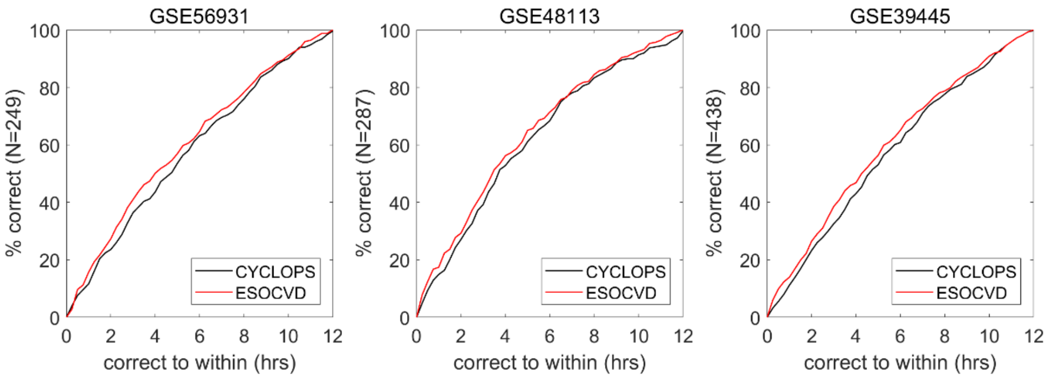


**Supplementary Fig. 13** The comparison between ESOCVD and CYCLOPS on three time-course datasets.

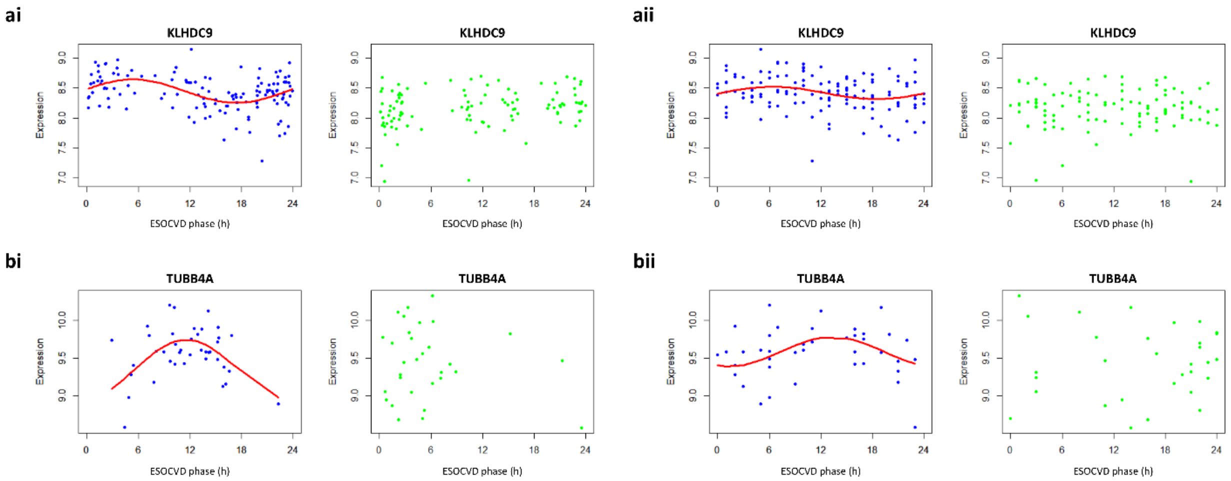


**Supplementary Fig. 14** Reconstructed expression profiles of two selected genes are plotted as a function of (i) CYCLOPS and (ii) ESOCVD.

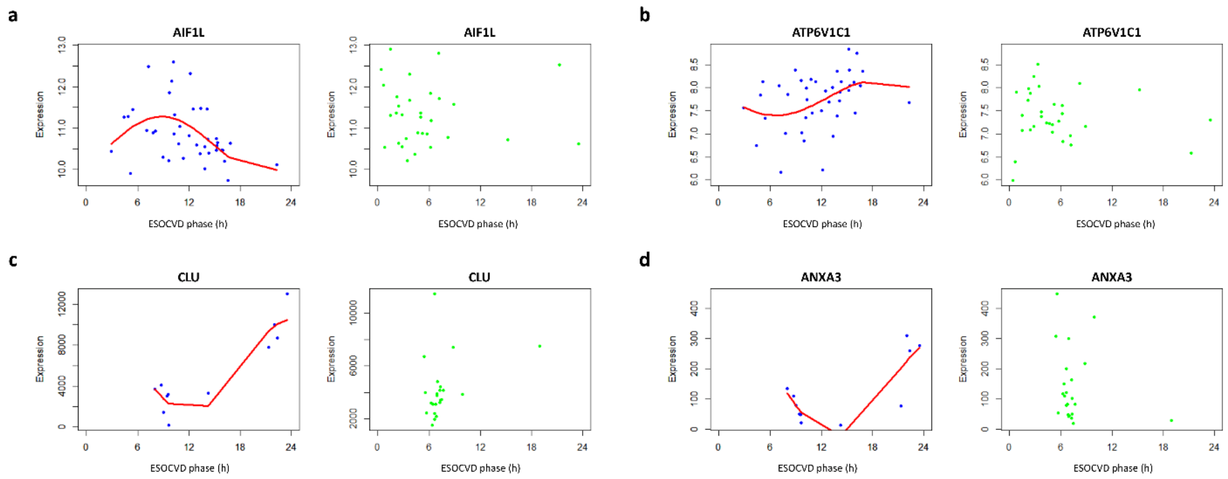


**Supplementary Fig 15** Reconstructed expression profiles of selected genes are plotted as a function of CYCLOPS phase.

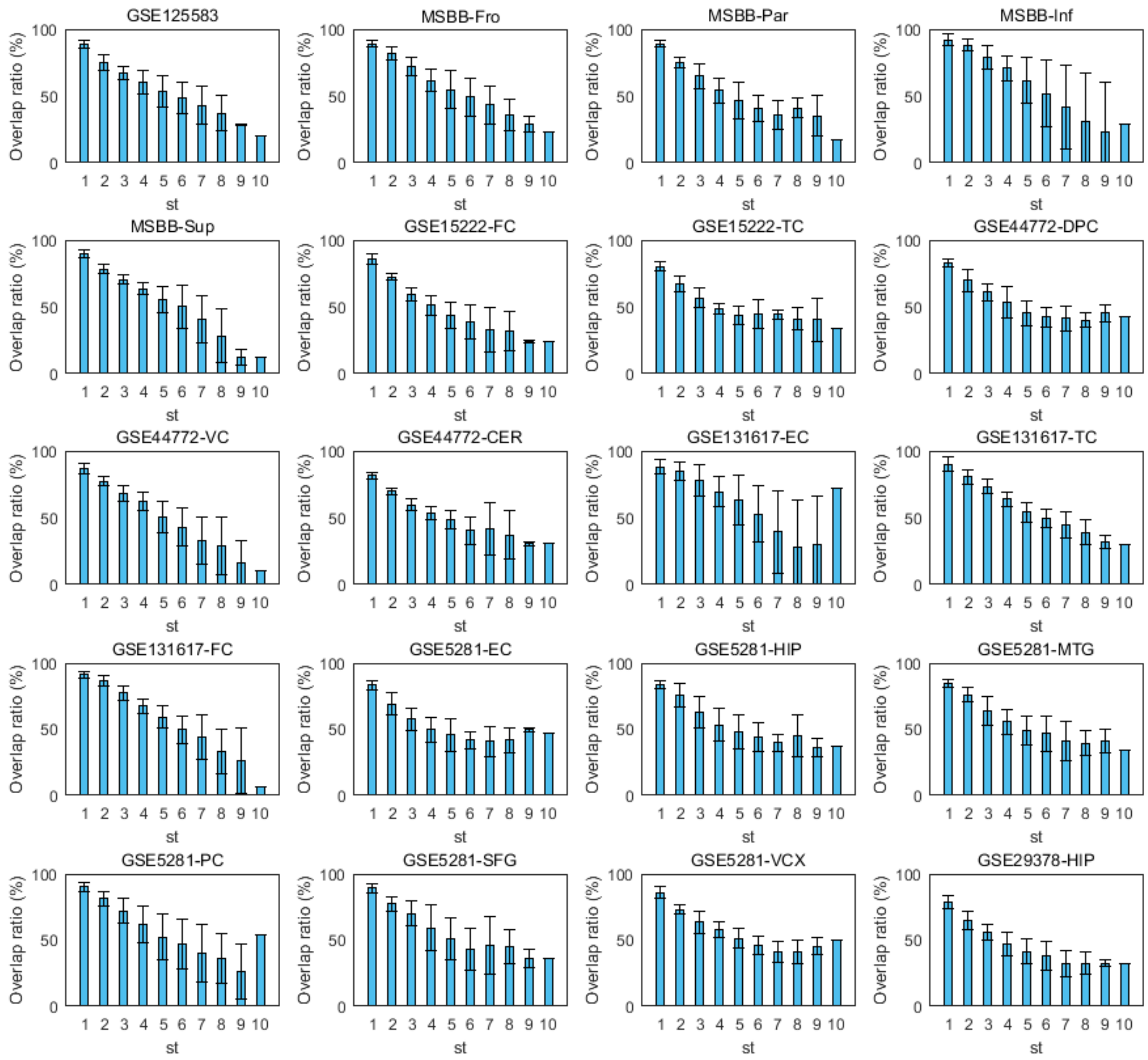


**Supplementary Fig. 16** The correlation between variable st and overlap ratio in heuristic search.

### Supplementary Tables

**Supplementary Table 1**. The information of all the gene expression profiles used in this study.

| Index | Data profiles | Dataset | Tissue | No. of samples | Control | Early | Mid | Late |
|---|---|---|---|---|---|---|---|---|
| 1 | GSE5281_EC | GSE5281 | EC | 23 | 13 | 0 | 0 | 10 |
| 2 | GSE5281_HIP | GSE5281 | HIP | 23 | 13 | 0 | 0 | 10 |
| 3 | GSE5281_MTG | GSE5281 | MTG | 28 | 12 | 0 | 0 | 16 |
| 4 | GSE5281_PC | GSE5281 | PC | 22 | 13 | 0 | 0 | 9 |
| 5 | GSE5281_SFG | GSE5281 | SFG | 34 | 11 | 0 | 0 | 23 |
| 6 | GSE5281_VCX | GSE5281 | VCX | 31 | 12 | 0 | 0 | 19 |
| 7 | GSE125583 | GSE125583 | FG | 289 | 70 | 0 | 0 | 219 |
| 8 | MSBB_Fro | MSBB | FP | 261 | 35 | 67 | 46 | 113 |
| 9 | MSBB_Par | MSBB | PG | 215 | 32 | 56 | 32 | 95 |
| 10 | MSBB_Inf | MSBB | IFG | 222 | 27 | 63 | 35 | 97 |
| 11 | MSBB_Sup | MSBB | STG | 240 | 33 | 58 | 37 | 112 |
| 12 | GSE15222_FC | GSE15222 | FC | 71 | 40 | 0 | 0 | 31 |
| 13 | GSE15222_TC | GSE15222 | TC | 241 | 135 | 0 | 0 | 106 |
| 14 | GSE44772_DPC | GSE44772 | DPC | 230 | 101 | 0 | 0 | 129 |
| 15 | GSE44772_VC | GSE44772 | VC | 230 | 101 | 0 | 0 | 129 |
| 16 | GSE44772_CER | GSE44772 | CER | 230 | 101 | 0 | 0 | 129 |
| 17 | GSE131617_FC | GSE131617 | FC* | 71 | 13 | 20 | 19 | 19 |
| 18 | GSE131617_TC | GSE131617 | TC* | 71 | 13 | 20 | 19 | 19 |
| 19 | GSE131617_EC | GSE131617 | EC* | 71 | 13 | 20 | 19 | 19 |
| 20 | GSE29378_HIP | GSE29378 | HIP | 72 | 37 | 0 | 0 | 35 |

**Supplementary Table 2**. The predicted and reference peak and trough of core clock genes.

| Clock genes | $Peak_p$ | $Trough_p$ | Regions (Dataset) | $Peak_r$ | $Trough_r$ | PMID |
|---|---|---|---|---|---|---|
| PER1 | 19 | 7 | MSBB_Sup | 18-20 | 6-8 | 20738730 |
| PER2 | 2 | 15 | GSE15222_FC | 2 | 14 | 17915197 |
| PER3 | 16 | 4 | MSBB_Sup | 16 | 4 | 20962856 |
| CRY1 | 19 | 7 | GSE131617_EC | 19 | 7 | 23896777 |
| CRY2 | 9 | 21 | GSE44772_VC | 9 | 21 | 11779462 |
| CLOCK | 3 | 17 | GSE131617_FC | 3 | 15 | 23896777 |
| NR1D1 | 19 | 7 | MSBB_Sup | 19 | 7 | 22629359 |
| NR1D2 | 22 | 8 | GSE5281_SFG | 22 | 10 | 23284293 |
| NPAS2 | 3 | 17 | GSE131617_FC | 3 | 15 | 23896777 |
| TEF | 9 | 20 | GSE131617_EC | 8 | 20 | 26576534 |

**Supplementary Table 3**. AD-related clock-controlled genes and inferred patterns of circadian variation.

| Gene name | Tissue | Dataset | Circadian pattern |
|---|---|---|---|
| ATXN10 | EC | GSE5281 | Gain of rhythmicity |
| SNCA | EC | GSE5281 | Gain of rhythmicity |
| BACE1 | EC | GSE5281 | Gain of rhythmicity |
| BACE2 | EC | GSE5281 | Gain of rhythmicity |
| ATXN7 | HIP | GSE5281 | Gain of rhythmicity |
| LRP1 | HIP | GSE5281 | Loss of rhythmicity |
| ATXN7 | MTG | GSE5281 | Gain of rhythmicity |
| ATXN10 | MTG | GSE5281 | Gain of rhythmicity |
| BACE1 | PC | GSE5281 | Loss of rhythmicity |
| ADAM10 | PC | GSE5281 | Loss of rhythmicity |
| SNCA | PC | GSE5281 | Gain of rhythmicity |
| ATXN10 | SFG | GSE5281 | Loss of rhythmicity |
| SNCA | SFG | GSE5281 | Loss of rhythmicity |
| SNCB | SFG | GSE5281 | Loss of rhythmicity |
| BACE1 | SFG | GSE5281 | Loss of rhythmicity |
| LRP1 | SFG | GSE5281 | Loss of rhythmicity |
| APH1A | VCX | GSE5281 | Gain of rhythmicity |
| SNCG | FG | GSE125583 | Gain of rhythmicity |
| ATXN2 | PG | MSBB | Gain of rhythmicity |
| SNCB | PG | MSBB | Gain of rhythmicity |
| LRP1 | PG | MSBB | Gain of rhythmicity |
| APBA1 | PG | MSBB | Gain of rhythmicity |
| ATXN2 | STG | MSBB | Loss of rhythmicity |
| SNCB | STG | MSBB | Loss of rhythmicity |
| BACE2 | STG | MSBB | Loss of rhythmicity |
| LRP1 | STG | MSBB | Loss of rhythmicity |
| APBA1 | STG | MSBB | Loss of rhythmicity |
| ATXN1 | FC | GSE15222 | Gain of rhythmicity |
| ATXN3 | FC | GSE15222 | Loss of rhythmicity |
| ATXN10 | FC | GSE15222 | Gain of rhythmicity |
| SNCA | FC | GSE15222 | Gain of rhythmicity |
| BACE2 | FC | GSE15222 | Gain of rhythmicity |
| APH1A | FC | GSE15222 | Loss of rhythmicity |
| LRP1 | FC | GSE15222 | Gain of rhythmicity |
| APOE | FC | GSE15222 | Gain of rhythmicity |
| BACE2 | TC | GSE15222 | Loss of rhythmicity |
| APP | TC | GSE15222 | Loss of rhythmicity |
| APOE | TC | GSE15222 | Loss of rhythmicity |
| ATXN10 | DPC | GSE44772 | Gain of rhythmicity |
| SNCA | DPC | GSE44772 | Loss of rhythmicity |
| SNCG | DPC | GSE44772 | Gain of rhythmicity |
| BACE2 | DPC | GSE44772 | Loss of rhythmicity |
| APBA1 | DPC | GSE44772 | Gain of rhythmicity |
| ATXN2 | VC | GSE44772 | Gain of rhythmicity |
| ATXN10 | VC | GSE44772 | Gain of rhythmicity |
| SNCA | VC | GSE44772 | Gain of rhythmicity |
| SNCB | VC | GSE44772 | Gain of rhythmicity |

| | | | |
|---|---|---|---|
| SNCG | VC | GSE44772 | Gain of rhythmicity |
| PSEN1 | VC | GSE44772 | Gain of rhythmicity |
| ADAM10 | VC | GSE44772 | Gain of rhythmicity |
| APH1A | VC | GSE44772 | Gain of rhythmicity |
| APBA1 | VC | GSE44772 | Gain of rhythmicity |
| IDE | VC | GSE44772 | Gain of rhythmicity |
| SNCG | CER | GSE44772 | Loss of rhythmicity |
| ATXN1 | FC | GSE131617 | Loss of rhythmicity |
| ATXN2 | Fc | GSE131617 | Loss of rhythmicity |
| ATXN3 | FC | GSE131617 | Loss of rhythmicity |
| ATXN7 | FC | GSE131617 | Loss of rhythmicity |
| IDE | FC | GSE131617 | Loss of rhythmicity |
| SNCB | TC | GSE131617 | Loss of rhythmicity |
| APH1A | EC | GSE131617 | Loss of rhythmicity |

**Supplementary Table 4**. Pathway level analysis of circadian changes with Superior frontal gyrus (SFG). KEGG, and GO biological processes sets that were overrepresented among those genes that change rhythmicity in SFG of AD patients are shown (P-value<0.05).

| **Pathway** | **P-value** |
|---|---|
| regulation of protein export from nucleus | 1.80E-04 |
| retrograde protein transport, ER to cytosol | 5.40E-04 |
| Ribosome | 8.10E-04 |
| microglial cell activation | 2.10E-03 |
| positive regulation of protein ubiquitination | 4.00E-03 |
| social behavior | 5.40E-03 |
| negative regulation of neuron apoptotic process | 8.60E-03 |
| Circadian rhythm | 9.90E-03 |
| proteasome-mediated ubiquitin-dependent protein catabolic process | 1.00E-02 |
| membrane fusion | 1.50E-02 |
| Other glycan degradation | 1.60E-02 |
| response to endoplasmic reticulum stress | 1.60E-02 |
| RNA splicing | 1.80E-02 |
| GTP metabolic process | 1.80E-02 |
| Estrogen signaling pathway | 2.00E-02 |
| Parkinson's disease | 2.10E-02 |
| chaperone-mediated protein complex assembly | 2.30E-02 |
| Oxytocin signaling pathway | 2.50E-02 |
| circadian regulation of gene expression | 2.50E-02 |
| Herpes simplex infection | 2.50E-02 |
| response to virus | 2.90E-02 |
| protein folding | 3.30E-02 |
| lysosome organization | 3.40E-02 |
| ER-associated ubiquitin-dependent protein catabolic process | 3.90E-02 |
| positive regulation of cell growth | 4.10E-02 |

**Supplementary Table 5**. Pathway level analysis of circadian changes with Superior temporal gyrus (STG). KEGG, and GO biological processes sets that were overrepresented among those genes that change rhythmicity in Sup of AD patients are shown (P-value<0.05).

| Pathway | P-value |
|---|---|
| Circadian rhythm | 2.40E-04 |
| circadian regulation of gene expression | 1.10E-03 |
| locomotory behavior | 5.35E-02 |
| cytoskeleton organization | 5.40E-03 |
| ABC transporters | 8.70E-03 |
| Axon guidance | 8.80E-03 |
| transferrin transport | 1.00E-02 |
| transcription, DNA-templated | 1.10E-02 |
| in utero embryonic development | 1.20E-02 |
| entrainment of circadian clock by photoperiod | 2.70E-02 |
| Bile secretion | 3.40E-02 |
| regulation of glucose import in response to insulin stimulus | 3.90E-02 |
| nuclear fragmentation involved in apoptotic nuclear change | 3.90E-02 |
| regulation of fatty acid transport | 3.90E-02 |
| nuclear inner membrane organization | 3.90E-02 |
| regulation of phosphatidylcholine catabolic process | 3.90E-02 |
| negative regulation of myeloid cell differentiation | 4.90E-02 |

**Supplementary Table 6**. Pathway level analysis of circadian changes with Frontal cortex (FC). KEGG, and GO biological processes sets that were overrepresented among those genes that change rhythmicity in FC of AD patients are shown (P-value<0.05).

| Pathway | P-value |
|---|---|
| Circadian rhythm | 2.70E-04 |
| Golgi organization | 1.00E-02 |
| ABC transporters | 1.00E-02 |
| transcription, DNA-templated | 1.80E-02 |
| fat cell differentiation | 3.40E-02 |
| mechanosensory behavior | 4.10E-02 |
| Circadian entrainment | 4.20E-02 |
| Glucagon signaling pathway | 4.30E-02 |

**Supplementary Table 7**. Pathway level analysis of circadian changes with Temporal cortex (TC). KEGG, and GO biological processes sets that were overrepresented among those genes that change rhythmicity in TC of AD patients are shown (P-value<0.05).

| Pathway | P-value |
|---|---|
| regulation of I-kappaB kinase/NF-kappaB signaling | 1.90E-03 |
| regulation of sodium ion transport | 5.50E-03 |
| cellular response to gamma radiation | 9.30E-03 |
| Endocrine and other factor-regulated calcium reabsorption | 1.80E-02 |
| Estrogen signaling pathway | 3.00E-02 |
| neuron apoptotic process | 3.30E-02 |
| Transcriptional misregulation in cancer | 3.90E-02 |
| receptor internalization | 4.30E-02 |
| response to hypoxia | 4.40E-02 |
| dynamin polymerization involved in mitochondrial fission | 4.50E-02 |
| Bile secretion | 4.70E-02 |

| | |
|---|---|
| endocytosis | 4.80E-02 |
| histone H3 acetylation | 4.90E-02 |

**Supplementary Table 8**. Pathway level analysis of circadian changes with Posterior cingulate (PC). KEGG, and GO biological processes sets that were overrepresented among those genes that change rhythmicity in PC of AD patients are shown (P-value<0.05).

| **Pathway** | **P-value** |
|---|---|
| Insulin signaling pathway | 4.20E-04 |
| regulation of cell shape | 2.20E-03 |
| Regulation of actin cytoskeleton | 1.80E-02 |
| negative regulation of epithelial cell proliferation | 2.30E-02 |
| positive regulation of ERK1 and ERK2 cascade | 3.00E-02 |
| Central carbon metabolism in cancer | 3.50E-02 |
| Pathways in cancer | 3.90E-02 |
| regulation of sequence-specific DNA binding transcription factor activity | 4.10E-02 |
| regulation of cell cycle | 4.90E-02 |

**Supplementary Table 9**. AD risk factors and inferred patterns of circadian variation.

| **Gene name** | **Tissue** | **Dataset** | **Circadian pattern** |
|---|---|---|---|
| CR1 | EC | GSE5281 | Gain of rhythmicity |
| PTK2B | HIP | GSE5281 | Gain of rhythmicity |
| ZCWPW1 | PC | GSE5281 | Loss of rhythmicity |
| MADD | PC | GSE5281 | Loss of rhythmicity |
| UNC5C | PC | GSE5281 | Loss of rhythmicity |
| CD2AP | SFG | GSE5281 | Loss of rhythmicity |
| PTK2B | SFG | GSE5281 | Loss of rhythmicity |
| PLD3 | SFG | GSE5281 | Loss of rhythmicity |
| MADD | SFG | GSE5281 | Loss of rhythmicity |
| CLU | SFG | GSE5281 | Loss of rhythmicity |
| MS4A6E | VCX | GSE5281 | Loss of rhythmicity |
| SORL1 | FG | GSE125583 | Gain of rhythmicity |
| TOMM40 | FG | GSE125583 | Gain of rhythmicity |
| CD33 | PG | MSBB | Loss of rhythmicity |
| CR1 | PG | MSBB | Gain of rhythmicity |
| NYAP1 | PG | MSBB | Gain of rhythmicity |
| MADD | PG | MSBB | Gain of rhythmicity |
| TREM2 | PG | MSBB | Gain of rhythmicity |
| NME8 | IFG | MSBB | Gain of rhythmicity |
| MS4A6A | IFG | MSBB | Gain of rhythmicity |
| BIN1 | STG | MSBB | Loss of rhythmicity |
| PTK2B | STG | MSBB | Loss of rhythmicity |
| PLD3 | STG | MSBB | Loss of rhythmicity |
| NYAP1 | STG | MSBB | Loss of rhythmicity |
| MADD | STG | MSBB | Loss of rhythmicity |
| MS4A6E | STG | MSBB | Loss of rhythmicity |
| PICALM | FC | GSE15222 | Gain of rhythmicity |
| MS4A6A | FC | GSE15222 | Gain of rhythmicity |
| MADD | FC | GSE15222 | Gain of rhythmicity |

| | | | |
|---|---|---|---|
| ABCB1 | FC | GSE15222 | Gain of rhythmicity |
| INPP5D | TC | GSE15222 | Loss of rhythmicity |
| PLD3 | DPC | GSE44772 | Gain of rhythmicity |
| ABCB1 | DPC | GSE44772 | Gain of rhythmicity |
| MS4A6A | DPC | GSE44772 | Loss of rhythmicity |
| RIN3 | DPC | GSE44772 | Loss of rhythmicity |
| BIN1 | VC | GSE44772 | Gain of rhythmicity |
| APP1 | VC | GSE44772 | Gain of rhythmicity |
| PLD3 | VC | GSE44772 | Gain of rhythmicity |
| NME8 | VC | GSE44772 | Gain of rhythmicity |
| ABCB1 | VC | GSE44772 | Gain of rhythmicity |
| UNC5C | VC | GSE44772 | Gain of rhythmicity |
| INPP5D | CER | GSE44772 | Gain of rhythmicity |
| PLD3 | CER | GSE44772 | Gain of rhythmicity |
| TREM2 | CER | GSE44772 | Loss of rhythmicity |
| ABCB1 | CER | GSE44772 | Loss of rhythmicity |
| ZCWPW1 | FC | GSE131617 | Loss of rhythmicity |
| CD2AP | FC | GSE131617 | Loss of rhythmicity |
| HLA-DRB5 | FC | GSE131617 | Gain of rhythmicity |
| ZW10 | EC | GSE131617 | Loss of rhythmicity |
| TOMM40 | EC | GSE131617 | Loss of rhythmicity |
| HLA-DRB5 | HIP | GSE29378 | Gain of rhythmicity |

**Supplementary Table 10**. The fold change values of the factors associated with CLU-related pathways in SFG. All the values were calculated with the data profile GSE5281_SFG.

| Gene name | Fold change (AD/CN) | Circadian pattern | Description |
|---|---|---|---|
| **JUN** | 1.529 | Loss of rhythmicity | Circadian gene; cofactor |
| **MAPK1** | 0.499 | Loss of rhythmicity | Circadian gene; cofactor |
| JUNB | 1.147 | | Circadian gene |
| **FOS** | **0.610** | Loss of rhythmicity | Circadian rhythm TF |
| **CLU** | **1.214** | Loss of rhythmicity | AD risk factor |

**Supplementary Table 11**. The fold change values of the factors associated with BIN1-related pathways in VC. All the values were calculated with the data profile GSE44772_VC.

| Gene name | Fold change (AD/CN) | Circadian pattern | Description |
|---|---|---|---|
| **BIN2** | **1.55** | | Circadian gene |
| **DNM2** | **1.389** | Gain of rhythmicity | Circadian gene; cofactor |
| WASL | 0.793 | Gain of rhythmicity | Circadian gene |
| DNM1 | 0.751 | | |
| AMPH | 0.589 | Gain of rhythmicity | Circadian gene |
| SYNJ1 | 0.683 | Gain of rhythmicity | Circadian gene |
| **BIN1** | **1.266** | Gain of rhythmicity | Circadian rhythm TF<br>AD risk factor |
| **MYC** | **1.387** | | Circadian gene |

**Supplementary Table 12**. The fold change values of the factors associated with BIN1-related pathways in STG. All the values were calculated with the data profile MSBB_Sup.

| Gene name | Fold change (AD/CN) | Circadian pattern | Description |
|---|---|---|---|
| **BIN2** | **1.172** | Loss of rhythmicity | Circadian gene; cofactor |
| DNM2 | 1.039 | Loss of rhythmicity | Circadian gene |
| WASL | 0.978 | | Circadian gene |
| DNM1 | 0.954 | | |
| AMPH | 0.965 | | Circadian gene |
| SYNJ1 | 0.959 | | Circadian gene |
| **BIN1** | **1.021** | Loss of rhythmicity | Circadian rhythmTF<br>AD risk factor |
| **MYC** | **1.156** | | Circadian gene |

**Supplementary Table 13**. The predicted and reference peak and trough of circadian genes.

| Gene name | $Peak_p$ | $Trough_p$ | Regions (Dataset) | Uniprot ID | $Peak_r$ | $Trough_r$ | CGDB ID |
|---|---|---|---|---|---|---|---|
| SDR39U1 | 16 | 3 | GSE5281_HIP | Q5M8N4 | 15 | 13 | CGD-MuM-014967 |
| NAA15 | 3 | 16 | GSE5281_PC | A0A0A6YW80 | 3 | 15 | CGD-MuM-010728 |
| SHISA9 | 17 | 3 | GSE5281_PC | Q9CZN4 | 15 | 3 | CGD-MuM-018139 |
| PHKA1 | 3 | 17 | GSE5281_PC | P18826 | 3 | 15 | CGD-MuM-013059 |
| FOS | 3 | 17 | GSE5281_SFG | P01101 | 16 | 8 | CGD-MuM-021107 |
| BCKDK | 16 | 3 | GSE5281_VCX | Q3UCB5 | 15 | 3 | CGD-MuM-014230 |
| NEURL2 | 14 | 2 | GSE125583 | A2A5J5 | 14 | 2 | CGD-MuM-011090 |
| ERF | 15 | 3 | GSE125583 | A0A0R4J0I0 | 15 | 3 | CGD-MuM-010910 |
| BHLHE22 | 6 | 17 | MSBB_Inf | A0A0R4J056 | 6 | 18 | CGD-MuM-010870 |
| FAM3A | 15 | 3 | MSBB_Sup | Q9D8T0 | 15 | 3 | CGD-MuM-018401 |
| PLA2G4E | 15 | 2 | MSBB_Sup | Q50L42 | 15 | 3 | CGD-MuM-014659 |
| CCT5 | 3 | 16 | GSE15222_FC | P80316 | 3 | 15 | CGD-MuM-013560 |
| KBTBD11 | 2 | 15 | GSE15222_FC | Q8BNW9 | 3 | 15 | CGD-MuM-016325 |
| HEXIM1 | 0 | 12 | GSE15222_TC | Q8R409 | 0 | 12 | CGD-MuM-017159 |
| AMD2 | 3 | 15 | GSE44772_DPC | D3Z6H8 | 3 | 15 | CGD-MuM-011781 |
| GLDC | 6 | 18 | GSE44772_VC | Q91W43 | 6 | 18 | CGD-MuM-017420 |
| PSMD5 | 0 | 12 | GSE44772_CER | Q8BJY1 | 0 | 12 | CGD-MuM-016238 |
| PHKA1 | 3 | 17 | GSE131617_FC | P18826 | 3 | 15 | CGD-MuM-013059 |
| DBT | 3 | 14 | GSE131617_FC | P53395 | 3 | 15 | CGD-MuM-013332 |
| AFF2 | 3 | 14 | GSE131617_FC | O55112 | 3 | 15 | CGD-MuM-012836 |
| METTL4 | 3 | 14 | GSE131617_FC | Q3U034 | 3 | 15 | CGD-MuM-014105 |
| KATNB1 | 17 | 3 | GSE131617_FC | Q8BG40 | 15 | 3 | CGD-MuM-016086 |
| C1QBP | 17 | 3 | GSE131617_FC | Q8R5L1 | 15 | 3 | CGD-MuM-017195 |
| ZMYM2 | 3 | 17 | GSE131617_FC | Q9CU65 | 3 | 15 | CGD-MuM-018025 |
| PPP1R13B | 3 | 17 | GSE131617_FC | Q62415 | 3 | 15 | CGD-MuM-015316 |
| SUGT1 | 3 | 17 | GSE131617_FC | Q9CX34 | 3 | 15 | CGD-MuM-018055 |
| PHF14 | 3 | 17 | GSE131617_FC | A0A0N4SV73 | 3 | 15 | CGD-MuM-010818 |
| XYLB | 3 | 17 | GSE131617_FC | Q3TNA1 | 3 | 15 | CGD-MuM-014000 |
| FAM3A | 17 | 3 | GSE131617_FC | Q9D8T0 | 15 | 3 | CGD-MuM-018401 |
| NFKBIB | 18 | 8 | GSE131617_FC | Q60778 | 18 | 6 | CGD-MuM-015144 |
| SMAD5 | 3 | 17 | GSE131617_FC | P97454 | 3 | 15 | CGD-MuM-013598 |
| ABCB7 | 3 | 17 | GSE131617_FC | Q61102 | 3 | 15 | CGD-MuM-015193 |

| | | | | | | | |
|---|---|---|---|---|---|---|---|
| RNF31 | 14 | 3 | GSE131617_FC | Q924T7 | 14 | 2 | CGD-MuM-017652 |
| PDLIM5 | 3 | 17 | GSE131617_FC | Q8CI51 | 3 | 15 | CGD-MuM-016812 |
| ZSCAN29 | 3 | 14 | GSE131617_FC | A2ARU5 | 2 | 14 | CGD-MuM-011238 |
| GALE | 17 | 3 | GSE131617_FC | Q8R059 | 15 | 3 | CGD-MuM-017042 |
| ENO3 | 17 | 3 | GSE131617_FC | Q5SX59 | 15 | 3 | CGD-MuM-015074 |
| INTS8 | 3 | 17 | GSE131617_FC | Q80V86 | 3 | 15 | CGD-MuM-015921 |
| CDC7 | 3 | 17 | GSE131617_FC | Q9Z0H0 | 3 | 15 | CGD-MuM-019079 |
| KLHDC9 | 17 | 3 | GSE131617_FC | Q3USL1 | 15 | 3 | CGD-MuM-014432 |
| WNT9A | 14 | 3 | GSE131617_FC | Q8R5M2 | 15 | 3 | CGD-MuM-017197 |
| IMPG2 | 3 | 17 | GSE131617_FC | Q80XH2 | 3 | 15 | CGD-MuM-015971 |
| GUCY2F | 4 | 3 | GSE131617_FC | Q5SDA5 | 15 | 3 | CGD-MuM-015009 |

**Supplementary Table 14** model convergence under different replicates (50, 100, and 200). Most of the patient samples involved in these datasets were collected from United States, except the patients in GES131617 were from Japan (see label "*").

| **Datasets** | **Data profiles** | **Brain regions** | **Likelihood (Repeat, 50)** | **Likelihood (Repeat, 100)** | **Likelihood (Repeat, 200)** |
|---|---|---|---|---|---|
| GSE125583 | GSE125583 | Fusiform gyrus | -1.0494e+04 | -1.0484e+04 | -1.0484e+04 |
| MSBB | MSBB_Fro | Frontal pole | -9.6228e+03 | -9.6113e+03 | -9.6113e+03 |
| | MSBB_Par | Parahippocampal gyrus | -7.8917e+03 | -7.8868e+03 | -7.8853e+03 |
| | MSBB_Inf | Inferior frontal gyrus | -8.2546e+03 | -8.2500e+03 | -8.2463e+03 |
| | MSBB_Sup | Superior temporal gyrus | -8.7360e+03 | -8.7324e+03 | -8.7320e+03 |
| GSE15222 | GSE15222_FC | Frontal cortex | -2.5823e+03 | -2.5813e+03 | -2.5813e+03 |
| | GSE15222_TC | Temporal cortex | -8.7479e+03 | -8.7456e+03 | -8.7453e+03 |
| GSE44772 | GSE44772_DPC | Dorsolateral Prefrontal cortex | -8.3589e+03 | -8.3541e+03 | -8.3520e+03 |
| | GSE44772_VC | Visual cortex | -8.2623e+03 | -8.2562e+03 | -8.2542e+03 |
| | GSE44772_CER | Cerebellum | -8.4804e+03 | -8.4694e+03 | -8.4694e+03 |
| GSE131617 | GSE131617_EC | Entorinal cortex* | -2.5105e+03 | -2.5078+e03 | -2.5073e+03 |
| | GSE131617_TC | Temporal cortex* | -2.4531e+03 | -2.4504e+03 | -2.4495e+03 |
| | GSE131617_FC | Frontal cortex* | -2.4914e+03 | -2.4877e+03 | -2.4859e+03 |
| GSE5281 | GSE5281_EC | Entorinal cortex | -813.67 | -811.93 | -811.75 |
| | GSE5281_HIP | Hippocampus | -830.31 | -828.73 | -828.61 |
| | GSE5281_MTG | Medial temporal gyrus | -1.0211e+03 | -1.0197e+03 | -1.0194e+03 |
| | GSE5281_PC | Posterior cingulate | -820.83 | -818.08 | -818.08 |
| | GSE5281_SFG | Superior frontal gyrus | -1.1939e+03 | -1.1921e+03 | -1.1912e+03 |
| | GSE5281_VCX | Primary visual cortex | -1.1267e+03 | -1.1249e+03 | -1.1247e+03 |
| GSE29378 | GSE29378_HIP | Hippocampus | -2.6356e+03 | -2.6332e+03 | -2.6331e+03 |
